# GraphCPLMQA: Assessing protein model quality based on deep graph coupled networks using protein language model

**DOI:** 10.1101/2023.05.16.540981

**Authors:** Dong Liu, Biao Zhang, Jun Liu, Hui Li, Le Song, Gui-Jun Zhang

## Abstract

Model quality evaluation is crucial part of protein structural biology. How to distinguish high-quality models from low-quality models, and to assess which high-quality models have relatively incorrect regions for improvement, are remain challenge. More importantly, the quality assessment of multimer models is a hot topic for structure predicton.In this work, we present GraphCPLMQA, a novel graph-coupled network that uses embeddings from protein language models to assess residue-level protein model quality. The GraphCPLMQA consists of a graph encoding module and a transform-based convolutional decoding module. In encoding module, the underlying relational representations of sequence and high-dimensional geometry structure are extracted by protein language models with Evolutionary Scale Modeling. In decoding module, the mapping connection between structure and quality are inferred by the representations and low-dimensional features. Specifically, the triangular location and residue level contact order features are designed to enhance the association between the local structure and the overall topology. Experimental results demonstrate that GraphCPLMQA using single-sequence embedding achieves the best performance compared to the CASP15 interface evaluation method in 9108 models of CASP15 multimer test set. In CAMEO blind test (2022-05-20∼2022-08-13), GraphCPLMQA ranked first compared to other servers. GraphCPLMQA also outperforms state-of-the-art methods on 19,035 models in CASP13 and CASP14 monomer test set. Finally, on AlphaFold2 datasets, GraphCPLMQA was superior to self-assessment of AlphaFold2 in MAE metric, and it was able to screen out better models than AlphaFold2.

## Introduction

Protein structure prediction plays an important role in biological research. In recent years, the development of deep learning has greatly advanced the transformation and progress of protein structure prediction. Many high-accuracy deep learning structure prediction methods have been developed, such as AlphaFold2 (1), RoseTTAFold (2), ESMFold (3), RGN2 (4) and PAthreader(5). More impressively, the collaboration between the European Molecular Biology Laboratory and DeepMind has predicted structures for over 200 millions proteins and made them freely available at the AlphaFold Protein Structure Database(31). In addition to experimental verification in the laboratory, there currently seems to be little prospect of designing computer programs that can independently validate AlphaFold2 models. Its own internal confidence estimates are likely to remain the best metrics of whether the model is correctly folded (6). This problem will become more significant when AlphaFold2 gets serious competition with other methods, such as ESMFold, OpenFold (7), which it will be inevitable. With the breakthrough of structure prediction, the reliability and usability of models are crucial parts, which are directly related to the efficiency of target discovery and drug design. Model quality assessment is important for structure prediction. Needless to say more, model quality assessment can further improve the accuracy of protein structure (8), and can also screen out the relatively best structure from multiple candidate models, which is critical for experimental scientists to analyze and verify.

Since CASP7, many methods for assessing the quality of protein models have been developed (19–21). In particular, single-model evaluation methods have received increasing attention and research, because they require only one model as input and show similar or better performance than consensus methods (22, 23). Features and networks are important for single-model quality assessment using deep learning. Features can explicitly describe the properties of proteins that include protein structural and non-structural features. For structural feature representation, some methods calculate inter-residue distances from atomic coordinates of protein models, and transform distances through spatial mapping to reflect the local structure and overall topology(9, 10). However, these methods only describe simple low-order distance relationships of protein geometric models and may ignore intrastructural connections in high-dimensional spaces. For nonstructural feature representations, the rosetta energy (24, 25) and statistical potential of the model represent the physicochemical information of the protein, such as ProQ3 (11) and VoroMQA(12). Particularly, sequence information implies the evolutionary relationship of proteins, which can improve the accuracy of model quality, as ProQ4 (13) and DeepAccNet-MSA (8). These methods just use sequence alignment information, it is more important to establish sequence-structure relationship. In addtion, for our in-house model quality assessment method, DeepUMQA (14) designed the residue-level USR (27) feature to characterize the topological relationship between the residuals and the overall structure. The improved version DeepUMQA2 (15) significantly improves the accuracy of model quality assessment by introducing co-evolution and template information, supplemented by an improved attention mechanism network framework. However, there is still space for improvement in network architecture.

Deep learning networks can capture potential connections within proteins. various neural networks that contain convolution, LSTM and graph networks, are used in model quality assessment methods, as ProQ3D (16), AngularQA (17) and GraphQA (18). These methods use a specific neural network architectures and build only one learning module. The learning mode of the network may be single, and the connection between the network architectures is not well utilized. Building blocks for specialized learning may help improve prediction accuracy. DeepAccNet utilizes 3D convolutional networks to obtain local atomic structure information, and then uses 2D convolutions to predict model quality. In addition, AlphaFold2 utilizes evolutionary blocks to encode sequence information and predict atomic coordinates and structural qulity in structural modules. Therefore, in model quality assessment, the network forms an encoder-decoder architecture, which can establish the connection among sequence, structure and quality to help improve the accuracy of model quality. The previous research studies show that structure, sequence, physicochemical information and deep learning network architecture are crucial for model quality assessment.

Protein language models are widely used in protein modeling and design tasks, which are trained unsupervised on protein databases to obtain embedding representations. In protein modeling tasks, sequence embeddings from protein language models are used to infer structural information, such as IgFold (28), ESMFold (3) and RGN2 (4). In sequence design tasks, structural embeddings from backbone atomic coordinates are used to predict protein sequences by networks, as ESM-IF1 (29). These methods show that protein language models establish a abundant connection between sequence and structure, which opens up the possibility of using language models in model quality assessment.

In this work, we propose GraphCPLMQA based on a deep graph coupled neural network framework using protein language models. Embeddings representations are generated by the protein language model ESM, which reflect sequence and structural properties. The embeddings that supplemented by structural features are input into a deep graph coupled network. The network consists of two parts: (1) The graph encoding network learns the latent connection between sequence and structure. (2) The transform-based convolutional decoding network obtains the mapping relationship between structure and quality to evaluate protein models. The results show that representations from language models and graph-coupled neural networks can learn the implicit relationshop among the sequence, structure and quality, which further improve the accuracy of model quality.

## Methods

In this section, we described the GraphCPLMQA method in three parts, including training and testing datasets, input features for proteins, and network architecture. In addition, we provide two versions according to different sequence embedding types, namely the full version of GraphCPLMQA (GraphCPLMQA-MSA) and the single sequence version of GraphCPLMQA (GraphCPLMQA-single). The pipeline was shown in Figure 1.

**Fig. 1.**
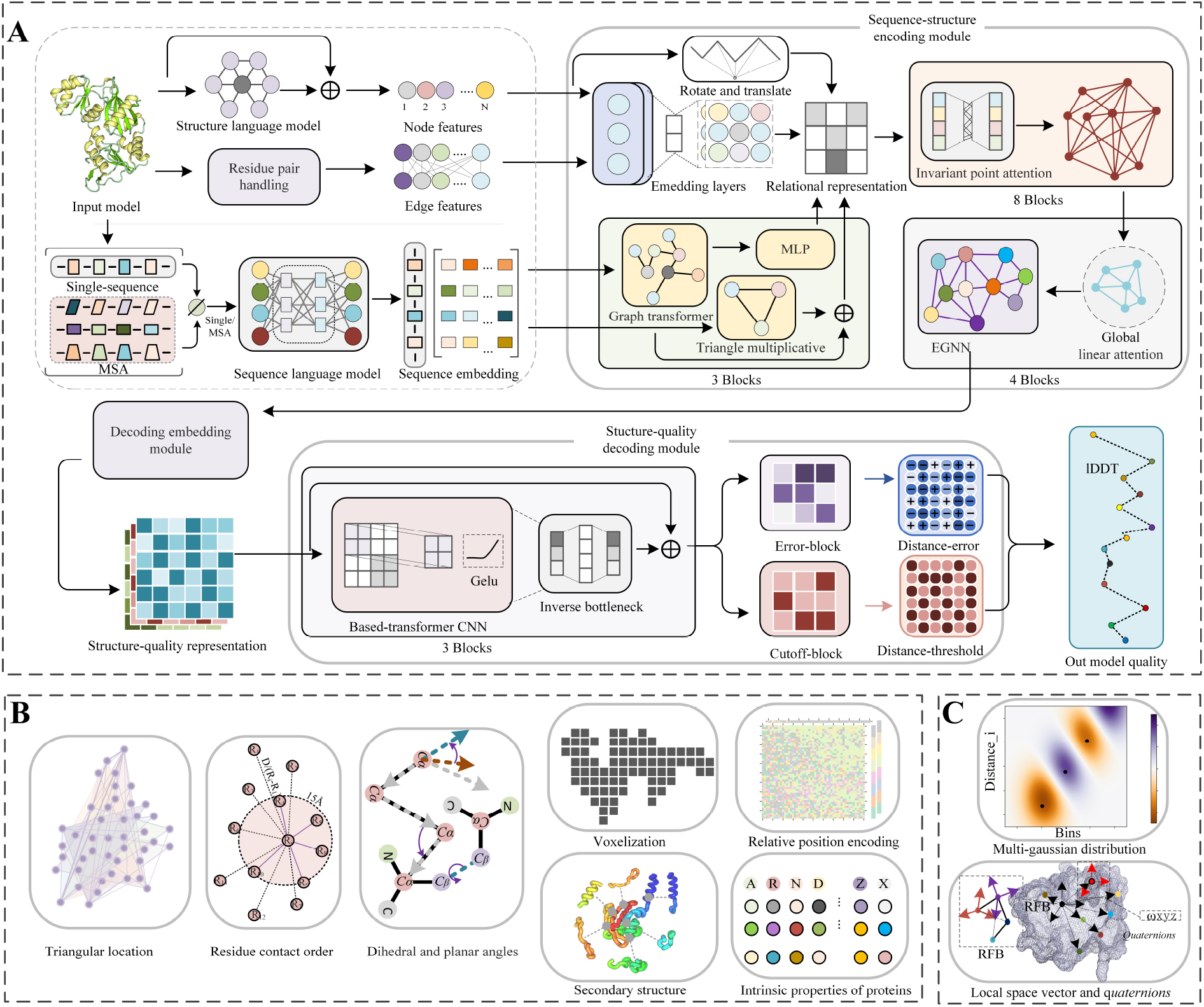
(A) The workflow of GraphCPLMQA. We extract the features in (B) and (C) along with the embedding representation from the protein structure where Single/MSA means that the input is single-sequence or MSA information corresponding to getting a single-sequence embedding or MSA embedding. In the Sequence-strucuture encoding module, we generate the relational representation of sequence and structure which inputs to the Structure-quality decoding module. Finally, the graph coupled network outputs the results of evaluating the model.

### A. Datasets

#### A.1. Dataset for training network model

The training dataset of GraphCPLMQA was constructed from Protein Data Bank (PDB) (30). A total of 15054 protiens were selected from the PDB (November 19, 2021) according to the following criterias: (1) minimum resolution <=2.5 Å, (2) protein length within 50-400 residues, (3) the sequence similarity of any to protiens less than 35%.

For each protein of the training set, three different approaches are used to generating decoys (structure models): structural dihedral adjustment, template modeling and deep learning-guided conformational changes(Supplement Figure S13). For structural dihedral adjustment, dihedral angles were fine-tuned on experimental structure (proteins from PDB). Each adjustment was followed by a fast relaxtion process. For template modeling, RosettaCM(32) and I-TASSER-MTD(33) were used to generate diverse structural models by utilize template structures with different accuracy and fragment libraries. For deep learning-guided conformational changes, our in-house method RocketX (57) was used to generated models by set different weights of geometric constraints. Additionally, the constrained conformations were refined to produce diverse models. Finally, after filtering for similar structures, a total of 1,378,676 protein models were obtained and utilized for training graph coupled network.

#### A.2. Test set construction

The performance of GraphCPLMQA is thoroughly tested on the model quality assessment datasets of CASP13, CASP14, and CASP15. GraphCPLMQA also participated in the continuous blind test of quality evaluation in CAMEO. All the test sets are constructed from the data provided by CASP and CAMEO official websites. The CASP13 and CASP14 test sets (CASP monomer test sets) were constructed by collecting models that were evaluated by all comparison methods and had a sequence similarity of less than 35% to the proteins in the training set. The CASP13 dataset collects 9390 structural models of 70 individual targets, and the CASP14 dataset collects 9645 models of 69 individual targets. The CASP15 test set (CASP multimer test set) was constructed by collecting structural models whose experimental structures had been released and length did not exceed 3000 residues, and collected 9108 models of 34 multimer targets. The CAMEO blind test set consists of 6 months CAMEO data (May 20, 2022 to August 13, 2022), which contains 1590 models of 189 proteins. Details of data can be found in Supplementary Tables S1-S3.

### B. Features

In the network, the input features can be classified into sequence and structural embeddings from language models, protein structural properties and protein physicochemical information. In graph networks, all features are divided into node and edge features, which are described in detail below.

#### B.1. Protein language embedding features

Most current model quality assessment methods use one-hot encoding for sequence embedding. However, there is sometimes problematic with only using one-hot encoding. One-hot encoding cannot represent the similarity or difference between amino acids effectively, as it may fail to capture the underlying relationships in protein sequences. Additionally, mapping sequences to structures using one-hot encoding can burden model quality assessment networks focused on structure learning. To characterize the structural information of protein models, some methods describe the relative positions of residues in a unified coordinate system, such as 3DCNN, Ornate, DeepAccNet, and DeepUMQA. However, these methods do not consider the implicit connections of residues in higher-dimensional spaces.

In this work, embeddings of protein sequences and geometric structures from ESM were employed to capture sequence-structure relationships. We have devised two distinct versions of the method to assess protein model quality, namely the GraphCPLMQA method, which utilizes the MSA language model, and the GraphCPLMQA-Single which employs a single sequence language model. For GraphCPLMQA-Single, the residue-level sequence embedding (1280-dim) of the query sequence is generated by ESM2(3) at the last layer (33rd) of the network. For GraphCPLMQA, the MSA of the input structure model is first produced through HHbits(37) searching against UniRef30 (51) and BFD(52), and then the searched MSA is fed into the ESM-MSA-1b(36) language model to derive residue-level sequence embeddings (768-dim) and row-attention embeddings between residues (144-dim) from the last layer (12th) of the network. Both versions of GraphCPLMQA utilize the ESM-1F1(29) to gengerate the structural embedding (512-dim) of the input backbone atomic coordinates.

#### B.2. Triangular location and residue level contact order

To describe the protein structure, the triangular location feature was designed, which is inspired by the residue-level USR from DeepUMQA (14). The feature characterizes the orientation and distance of the local structure in the overall topology. To construct the triangular location feature, the farthest point 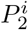 was identified for the *C*_*α*_ coordinate of residue 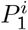 in the protein structure, and 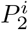 was taken as the center to find the farthest point 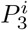 (excluding 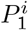). These three points formed a triangle *S*_*i*_, with *N* (number of residues) small triangles outlining the fundamental shape of the protein structure. In each triangle *S*_*i*_, the side lengths were 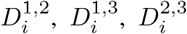, and the average distance from all residues to these three points were calculated as 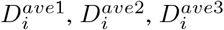. Finally, a local coordinate system Γ_*i*_ was constructed with 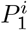 to characterize the position of the triangle in space. The local coordinate system described the orientation of the local structure in the overall topology. The calculation process is as follows:

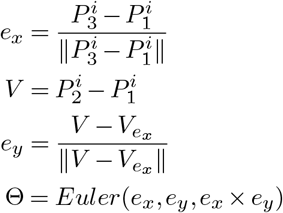

where 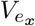 denotes that the projection length of *V* on *e*_*x*_ multiplied by *e*_*x*_ to obtain a projection vector. In the above, *Euler* is the mapping function from the local coordinate system to the Euler angles.

Contact order (50) is used to describe the overall topology complexity. We further extend to residue-level features to describe the complexity of local structures and between local structures. The calculation process is as follows:

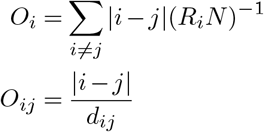

where *i, j* are index of residues; *R*_*i*_ is the number of adjacent residues within a distance of 15 Å for residue *i*; *d*_*ij*_ is the distance between residue *i* and residue *j*; *N* is protein length.

#### B.3. Protein node features and edge features

The positional order and properties of amino acids are critical to protein structure. The relative position encoding method in Tansformer (49) was employed to encode the sequence order, and take the sine and cosine of the position differences between the sequences as node fetures of residues. The properties of amino acids are represented by Meiler (38) and Blosum62 (39). To characterize the information of the secondary structure, DSSP (40) was used. The voxelization(10) of protein structures with rotation-translational invariance further complements the overall topological information. To capture the spatial arrangement of residues within the protein structure, vectors between backbone atoms are used to represent dihedral and plane angles.

Local vector 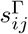 is used to describe the relative positional relationship of residues, and the rotation transformation *Q*_*ij*_ represent the relationship between each local spatial structure(See Supplementary Text S4 for details). In order to map the distances from the main chain atoms to a high-dimensional space, different interval Gaussian functions are employed to disperse the distances. Additionlly, distance map features are computed between the *C*_*β*_ atoms and the tip atoms (8), which complement the edge information of the graph network. The inter-residue Rosetta energy terms are used to represent the physicochemical information of the protein. The detailed dimension information of all features is in the Supplementary Table S5.

### C. Network architecture

The graph coupled network of GraphCPLMQA consists of two parts. One encodes sequence and structure information, and the other decodes structure and quality relationships. The architecture and training process of the coupled network are described below. The network parameters are described in detail in the Supplementary Table S6.

#### C.1. Sequence-structure encoding module

In encoding module, a protein graph is typically represented as 𝒢 = (𝒱, *ℰ, 𝒳*). In 𝒢 protein graph, 𝒱 = {*υ*_1_, *υ*_2_, …, *υ*_*N*_} is the set of residue, ℰ = {*ε*_*ij*_}_*i*≠*j*_ is the set of edge between residues, where each 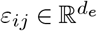 is the feature vector between residue *i* and residue *j*. 𝒳 = {*χ*_*i*_ ∈ *ℝ*^3*×*5^} represents the *C, O, N, C*_*α*_, *C*_*β*_ coordinates of the backbone atom for residue *i*.

The residual embedding 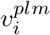 and residual attention 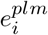 from the protein language model were fed into encoding module. Then, the embedding features are updated by a graph transformer (GT) (41) and triangular multiplication block(1), which can delve deeper into spatial geometric information from the abundant relationship between sequence and structure. In the GT layer, the multi-head attention of each node *i* with all other nodes *j* and the new nodes 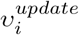 shows as Supplementary Text S2, S5.

The triangular multiplication operation in AlphaFold2 is used to update the edges between residues, along with the invariant point attention module to establish the intrinsic relationship between protein sequence and structure in high-dimensional space. By incorporating the sequence information into the model, we update the rotation and translation of the process. Finally, a geometric spatial constraint is obtained that is strongly associated with the sequence.

Equivalent graph neural network (EGNN) (42) further updates the node information, which helps to correct the unreasonable spatial constraints on the nodes in preparation for input decoding. In the EGNN architecture, residue node *i* is not globally connected to all other residue nodes. To form a new graph 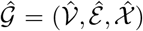, we search for the K nearest residue nodes in Euclidean space for each node *i*. The global linear attention is initially utilized to update the structure node, which follows a similar implementation to the aforementioned process. Following this, a graph-equivariant update method is used for residues as Supplementary Text S3. The proof process of invariance is in the Supplementary Text S1.

#### C.2. Structure-quality decoding module

In the decoding embedding module, we extract the node representation 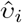 and the edge representation 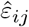 from the output of the encoding module. Moreover, the embedding and structural features are regenerated by a new network function *W* ^*^ for new nodes 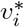 and edges 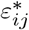 where the function utilizes new parameters. These all features are combined to generate a Structure-Quality representation as follows:

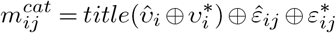

where ⨁ denotes the concatenation of feature vectors; *tile* indicates horizontal striping of node features into edge features.

In the Structure-quality decoding module, a residual network based on a transform strategy is employed, which consists of main residual blocks and branch residual blocks (Error-Block and Cutoff-Block). Each residual block comprises three 2-dimensional convolutional layers with different expansion rate coefficients and a normalization operation. We take the GELU(43) activation function and inverted bottleneck method which is one important design in every transformer block (44). Moreover, the convolutional network layer is added in the residual block of the branch to improve the prediction of distance error and threshold, as follows:

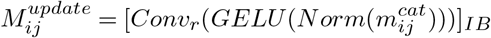

where *Conv*_*r*_ are 2-dim convolutional networks with different dilation coefficients *r* = *p*^2^(*p* = 1, …, 4); *IB* is the operations of inverted bottleneck.

The distance-error 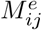 and the distance-threshold 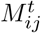 are obtained from the base-transformer residual network. 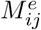 is the predicted distance error between the real structure and the model structure, and the distance threshold 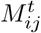 is the distance value within 15 Å where the threshold range is from lDDT (45). Finally, we calculate the local quality score as Supplementary Text S6.

#### C.3. Training procedure

The graph-coupled network model is trained using a combination of model quality and geometric constraint loss terms. To improve the efficacy of the sequence-structure encoding module, we use actual geometric structure information to constrain the encoding output by the mean square error between atomic coordinates and the L1 loss function. This approach helps in decoding the underlying information of structure and quality. In the decoding module, we compute the cross-entropy loss for the distance error and threshold, where the loss term for the threshold is the binary entropy. Finally, the loss function of the model quality is the mean squared error (Supplementary Text S7). In addition, to preserve the model during training, only 4% of the dataset structure is used for validation. For optimization, we utilized the AdamW (46) optimizer with a learning rate of 0.001, which decays at a rate of 1%. The top five models were trained with a batch size of one protein model for 100 epochs, which took approximately 120 hours on a single A100 GPU.

## Results and discussion

We use the constructed structure dataset to train the graph coupled network, which is used to test the non-redundant CASP proteins. Morever, We participated in the blind test of CAMEO and analyzed the quality assessment data. During the test, the global quality assessment (Global QA) and the accuracy of the local structure quality (Local QA) were used. Local QA describes the quality of each residue, where lDDT is used to evaluate the residue quality. Global QA describes the overall quality of the protein model structure by calculating the mean value of the Local QA. Pearson, Kendall (53), AUC (54), Mean absolute error (MAE), Mean Squared Error (MSE), and Top1loss (56) are commonly used evaluation metrics for Global QA. Similarly, Pearson, Spearman (55), Kendall, AUC, MAE, and MSE are used as evaluation metrics for Local QA. Pearson estimated the correlation between the predicted and real qulity of local residues or overall structure. Greater values indicate a stronger correlation and improved performance of the method. The error between the predicted quality and the real quality was measured using MAE and MSE, with the magnitude of the value indicating the gap from the real quality. A smaller value indicates better performance in predicting the quality. These metrics help assess the performance of models in terms of their accuracy and ability to make predictions.

### D. Results on the recent CASP15 multimer test set

With the precision breakthrough of Alphafold2 in monomers, research into multimers has become a top priority. Similarly, assessing the quality of multimer interfaces is a future frontier and presents a challenge. The lack of effective MSA information greatly increases the difficulty of predicting and evaluating multimers. However, GraphCPLMQA-Single employs a single-sequence embedding to assess interface quality, as shown in Figure 2 and Supplementary Table S14, S15.

**Fig. 2.**
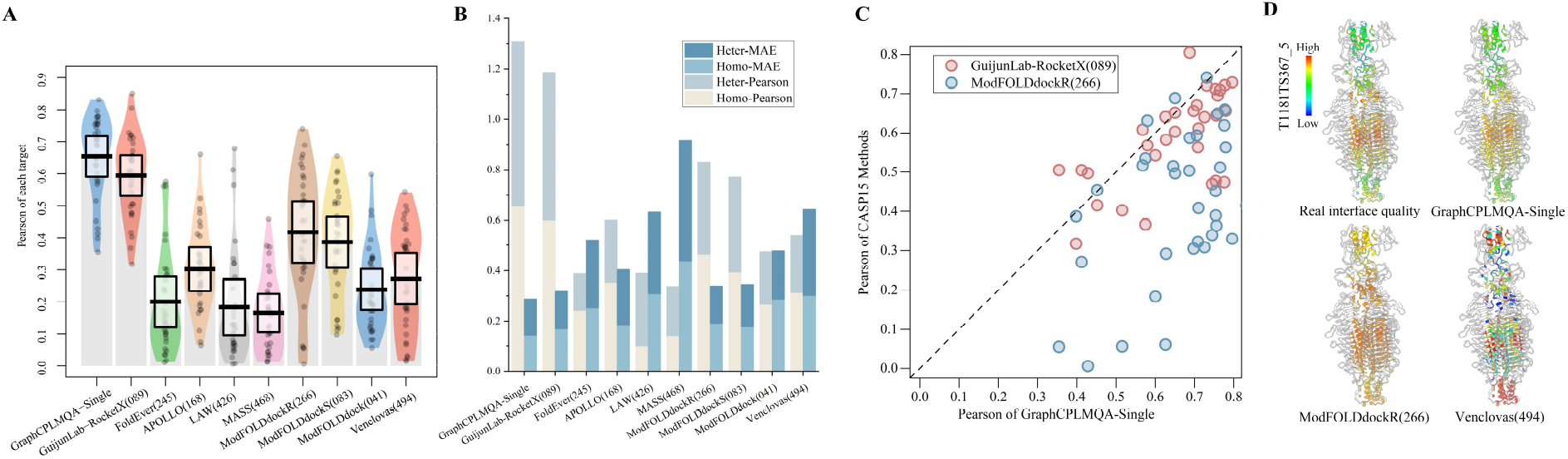
Test results for interface residues in the CASP multimer test (CASP15). (A) The pirate graph shows the Pearson correlation of different methods in predicting the quality of the multimer interface and the quality of the real multimer interface, where the horizontal line is the mean line. (B) The histogram depicts the performance analysis of different methods on CASP15 homo-oligomers and hetero-oligomers. (C) The scatterplot shows GraphCPLMQA-Single compared with the top method GuijunLab-RocketX and the second method ModFOLDdockr in recent CASP15 interface quality evaluation.(D) For model T1181_TS367_5, different methods predict the quality distribution at the multimer interface.

GraphCPLMQA-Single is compared with 9 methods in CASP15 for predicting multimer interfaces. On Pearson metrics, GraphCPLMQA-Single has improved by 23.6% compared to ModFOLDdockR(266), and its interface quality prediction ranks second in CASP15(Supplementary Figure S14 D). On the MAE metric, we observed that our method outperforms GuijunLab-RocketX(089), which is considered one of the top-performing methods for CASP15 multimer evaluation, in terms of predicting interface quality(Supplementary Figure S14 A). In other metrics, our method also achieves the highest performance with Spearman(0.617), Kendall(0.45), AUC(0.844), MSE(0.035), MAE(0.144), compared with other methods. For each target, our method predicts results with higher stability and accuracy than other methods (Figure 2 A, Supplementary Figure S14 B).

In particular, we compare with ModFOLDdockr (266) on each target. The results show that prediction accuracy of GraphCPLMQA-Single outperforms ModFOLDdockr (Figure 2C, Supplementary Figure S14 C). Evaluation of the model interface for T1181(Figure 2D), the predicted quality of GraphCPLMQA-Single is closer to the real quality, where the quality corresponds to the change of color (low: blue, high: red). Further, we analyzed the performance of the evaluation method on different types of multimeres (homo-oligomers and hetero-oligomers) as shown Figure 2B, Supplementary Table S14. Interestingly, the performance of our method on different types of multimeres is basically consistent. GraphCPLMQA-Single remains at the highest accuracy for evaluating interfaces in both homo-oligomers and hetero-oligomers.

The above results show that the performance of GraphCPLMQA-Single surpasses other CASP15 methods. Although GraphCPLMQA-Single is trained on monomer data, it performs well on multimer interface evaluation. GraphCPLMQA-Single shows potential for extension to evaluate multimer interfaces, which may be attributed to the following reasons. Firstly, the network has learned the evaluation mode of the local structural quality on proteins; secondly, the network takes the input protein structure as a whole, regardless of whether it is a multimer or a monomer; finally, the features of the network can be described structural and sequence information of the multimer. However, our method still has deficiencies in the interface evaluation of the CASP15 multimer test set. It can be seen from Supplementary Figure 14 E, F that the accuracy of both ends of the abscissa is relatively low, which is arranged from short to long according to the length of the target, and the accuracy of the middle part is relatively high and stable. This shows that the length of the multimer model will have a certain degree of impact on the evaluation accuracy of GraphCPLMQA-Single.

### E. Results on the CASP monomer test set

For 9390 protein models of CASP13 and 9645 protein models of CASP14, GraphCPLMQA and GraphCPLMQA-Single compare performance with state-of-the-art methods on the CASP monomer dataset (Figure 3, Supplementary Figure S9, Table S7, S8). In the monomer test set, GraphCPLMQA achieves the highest accuracy on both Global QA and Local QA metrics, surpassing other comparable methods. QMEANDisCo and DeepAcc-MSA were one of the best-performing model quality assessment methods in CASP13 and CASP14, respectively. GraphCPLMQA is analyzed using Pearson and MAE with QMEANDisCo and DeepAcc-MSA for residues of all models. In terms of global quality, GraphCPLMQA had a Pearson of 0.927, representing a 3% improvement over DeepAcc-MSA (Supplementary Figure S9 A-C). We analyzed the quality distribution of the T1042 and T0962 models, and the results predicted by GraphCPLMQA are basically the same as the real quality, where the range of color is the reference real quality distribution (Supplementary Figure S9 D, E). There are other models of case in Supplementary Figure S10. GraphCPLMQA-Single uses a single-sequence embedding for comparison with methods that use single-sequence information. Similarly, GraphCPLMQA-Single outperforms other methods using single-sequence information on all metrics, such as DeepUMQA, QMEANDisCo, ModFOLD8, DeepAcc, etc. GraphCPLMQA-Single shows similar performance to DeepAcc-MSA using MSA. In addition, the scatterplots of GraphCPLMQA and GraphCPLMQA-Single compared with other methods in Supplementary Figure S2-S8. The above results show that embeddings from language models and graph-coupled networks improve the accuracy of model quality assessment. The impact of different parts on the accuracy of the method can be seen in the Ablation studies.

**Fig. 3.**
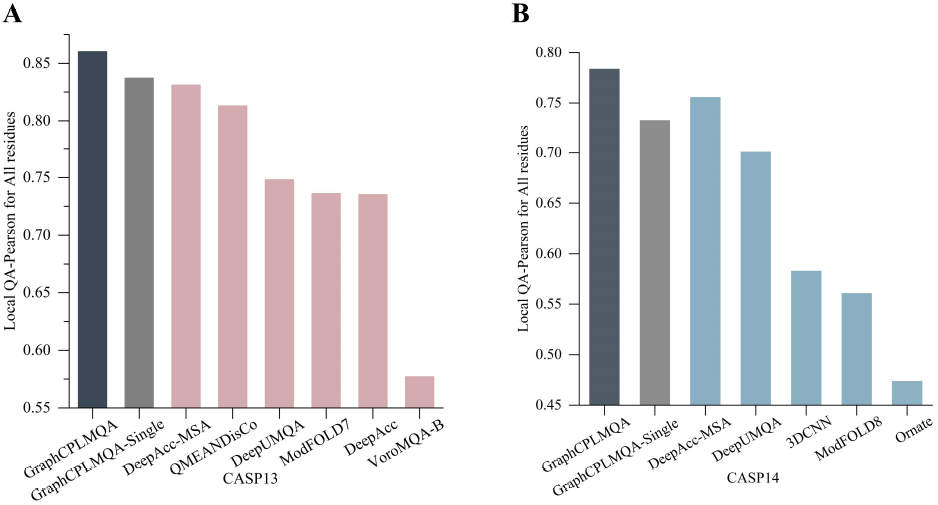
Result on the CASP monomer test set. (A),(B) for the all residue of CASP13 and CASP14 test set, GraphCPLMQA and GraphCPLMQA-Single were compared with other methods based on the Pearson correlation between the predicted and real quality of residues.

### F. Results on the CAMEO blind test

We developed the server ZJUT-GraphCPLMQA (server 46) based on the GraphCPLMQA method to participate in CAEMO-QE. Additionally, more than 3134 protein models have been evaluated so far. In the competition, other participating servers included DeepUMQA2, DeepUMQA, QMEANDisCo3, ProQ3D_LDDT, VoroMQA_v2, QMEAN 3, ProQ2, ModFOLD8, ProQ3D, ModFOLD6, VoroMQA_sw5. We download the test data on the CAMEO official website (from May 20, 2022 to August 13, 2022). Among the 128,018 residues in the CAEMO blind test, the evaluation accuracies of GraphCPLMQA for local residuals are Pearson (0.891), Kendall (0.680), AUC (0.942), and MAE (0.081), all of which exceed the accuracies of other servers, and MSE (0.015) is inferior to DeepUMQA2 (Table 1). The local Kendall distributions

**Table 1.** Results of ZJUT-GraphCPLMQA (server 46) on CAMEO test set

| Methods | Local QA |  |  | Global QA |  |  |
| --- | --- | --- | --- | --- | --- | --- |
|  | Pearson | AUC | MAE | Pearson | AUC | Top1loss |
| GraphCPLMQA<sup>‡</sup> | **0.891** | **0.942** | **0.081** | **0.924** | **0.967** | **0.007** |
| DeepUMQA2 | 0.878 | 0.941 | 0.083 | 0.908 | 0.966 | 0.016 |
| DeepUMQA | 0.835 | 0.923 | 0.099 | 0.874 | 0.956 | 0.015 |
| QMEANDisCo3 | 0.829 | 0.925 | / | 0.885 | 0.962 | 0.035 |
| ModFOLD8 | 0.771 | 0.894 | 0.127 | 0.824 | 0.920 | 0.066 |
| ProQ3D_LDDT | 0.769 | 0.887 | 0.151 | 0.824 | 0.904 | 0.024 |
| ProQ3 | 0.749 | 0.881 | 0.156 | 0.825 | 0.914 | 0.028 |
| VoroMQA_v2 | 0.733 | 0.880 | 0.176 | 0.824 | 0.926 | 0.015 |
| ProQ2 | 0.717 | 0.859 | 0.176 | 0.812 | 0.900 | 0.035 |
| ModFOLD6 | 0.716 | 0.877 | 0.182 | 0.811 | 0.906 | 0.077 |
| ProQ3D | 0.708 | 0.853 | 0.151 | 0.811 | 0.892 | 0.032 |
| QMEAN3 | 0.705 | 0.886 | 0.333 | 0.807 | 0.921 | 0.021 |
| VoroMQ_sw5 | 0.596 | 0.828 | 0.188 | 0.766 | 0.878 | 0.023 |
| **Note:** GraphCPLMQA<sup>‡</sup> enters CAEMO as group ZJUT-GraphCPLMQA (server 46). |  |  |  |  |  |  |

Methods Local QA Global QA Pearson AUC MAE Pearson AUC Top1loss of target proteins are shown in Supplementary Figure S16 D. In the global quality assessment, the Pearson and AUC accuracy of GraphCPLMQA are 0.924 and 0.967, higher than QMEANDisCo 3, and the accuracy of MAE (0.077) is second only to DeepUMQA2. An analysis was conducted on the quality predictions of different servers for the protein model 8D1X_D_20_1. The quality trend predicted by the server and the real quality trend are shown in Supplementary Figure S16 E. Further, in Supplementary Figure S16 F∼I, the model quality corresponds to the change of color, and it can be clearly seen that the accuracy of our prediction is higher. There are other models of case in Supplementary Figure S11, and blind test results in the Supplementary Figure S1.

### G. Ablation studies

The impact of the features and network architecture for GraphCPLMQA and GraphCPLMQA-Single on the CASP monomer test datasets was analyzed (Figure 4, Supplementary Figure S15). At the sequence feature level, we compare the performance of GraphCPLMQA using MSA embedding with GraphCPLMQA-single using single sequence embedding. In terms of various evaluation metrics, the performance of GraphCPLMQA with MSA embedding was superior to that of the counterpart without MSA (Figure 4). This suggests that the MSA contains richer structural information compared to single sequence. The embeddings derived from MSA provide better guidance for evaluating model quality. Further, we analysis in detail the impact of components on method performance below.

**Fig. 4.**
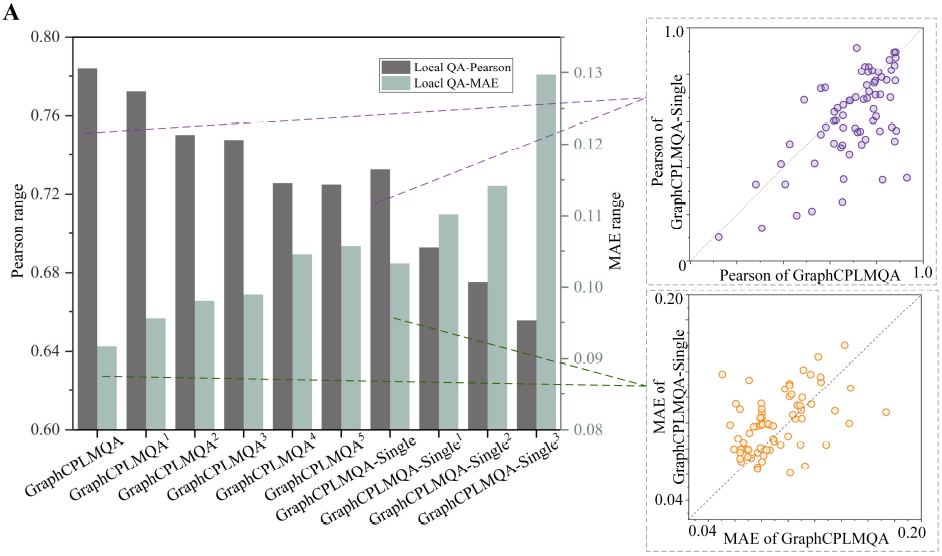
The impact of various components on the performance of GraphCPLMQA and GraphCPLMQA-Single.

To investigate the impact of the components, we modified the full version of GraphCPLMQA by removing some features and changing the network architecture. Different network models were retrained to test the results and analyze the effect by the following modified part: (1) GraphCPLMQA^1^ was replacing the transformer strategy inverted bottleneck and GELU (43) with residual block and ReLU (47), (2) for GraphCPLMQA^2^, the structural embeddings were removed from the language model in GraphCPLMQA^1^, (3) output mode of the encoding module(GraphCPLMQA^3^) and the connection architecture between modules(GraphCPLMQA^4^) on the GraphCPLMQA^2^ model was changed. (4) Based on GraphCPLMQA^2^model, the triangle position and residual-level contact order features were removed(GraphCPLMQA^5^). These components have an important impact on the performance of GraphCPLMQA (See the Supplementary Text S8 for details).

On GraphCPLMQA-Single model, we analyze the effect of different single sequence language models on the performance of the method. Specifically, the network models GraphCPLMQA-Single^1^ and GraphCPLMQA-Single^2^ were retrained with the high-dimensional sequence embedding of the ESM-1b(34) and ESM-1v(35) language models, respectively. The use of these embeddings resulted in a noteworthy decrease in performance. Furthermore, the input pattern of sequence embedding was explored using the GraphCPLMQA-Single^3^ network model. Although we acquired information from all embedding layers by averaging the sequence embeddings obtained from all network layers of the ESM-1v language model, this approach introduced significant noise that could potentially impact the accuracy of predictions.

### H. Compared to AlphaFold2

Since the results of AlphaFold2 prediction in CASP14 have no self-assessment pLDDT, we used the official website code of AlphaFold2 (https://github.com/deepmind/alphafold) to predict 69 sequences of CASP14. AlphaFold2 produced five output models for each sequence, resulting in a total of 345 models with pLDDT. GraphCPLMQA and GraphCPLMQA-Single assessed each model quality of AlphaFold2, respectively. The quality of GraphCPLMQA assessment exceeds self-assessment of AlphaFold2 on MAE (Figure 5A, Supplementary Figure S12). On 345 AlphaFold2 models, 253 evaluated results exceeded AlphaFold2 pLDDT. On the 207 AlphaFold2 structures without template information, GraphCPLMQA had 150 better evaluated results. GraphCPLMQA-Single performed slightly better than AlphaFold2 pLDDT on all structures, including those without a template. Furthermore, we analyze the evaluation of GraphCPLMQA on Alphafold medium and high precision models versus its self-evaluation as Supplementary Figure S17. The results show that our method helps to complement the deficiencies that exist in the pLDDT of AlphaFold2. In future studies, our method may also provide a valuable reference for predicting models in AlphaFoldDB that do not have native structures.

**Fig. 5.**
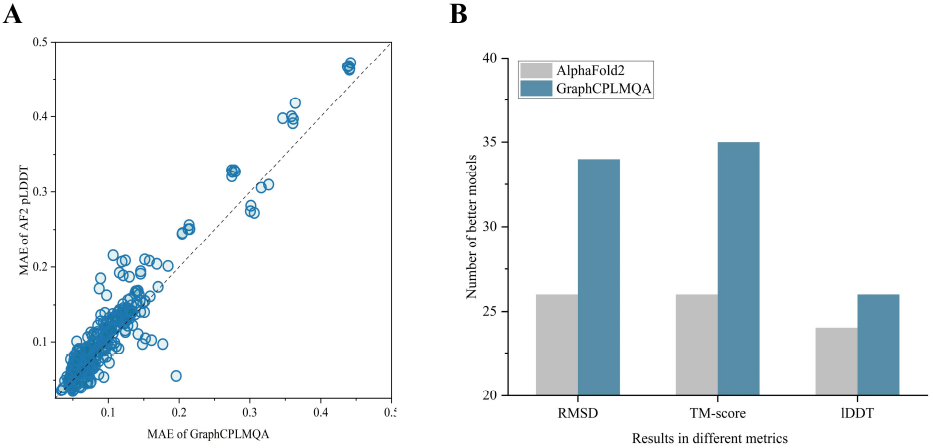
Results of GraphCPLMQA evaluation of AlphaFold2 structures compared to the AlphaFold2 pLDDT self-assessment. (A) The scatterplot further shows the distribution on the MAE. (B) GraphCPLMQA and AlphaFold2 select better models on different metrics.

In addition, the ability to select AlphaFold2 models was also evaluated (Figure 5B). Specifically, for AlphaFold2 dataset, the best structure was selected from the five predicted models of AlphaFold2 and compared to the best predicted model (rank_0) of AlphaFold2. On the test set, the average selection error of GraphCPLMQA was lower than that of AlphaFold2 in RMSD (0.672), TM-score(48) (0.015) and lDDT (0.008). Furthermore, we had 34, 35 and 26 structures in RMSD, TM-score, lDDT better than the best structure of AlphaFold2. These results demonstrate that GraphCPLMQA outperformed self-evaluation of AlphaFold2 in selecting the best model.

## Conclusion

In this study, we propose GraphCPLMQA, a novel approach for evaluating model quality that combines graph coupled networks and embeddings from protein language models. GraphCPLMQA utilizes sequences and structure embeddings, as well as additional model features, to establish the relationship among sequence, structure and quality. By predicting protein model quality scores, GraphCPLMQA outperform other state-of-the-art assessment methods in terms of accuracy on the CASP13, CASP14, CASP15 and CAEMO test sets. GraphCPLMQA also achieves excellent results in the continuous evaluation of CAMEO-QE. GraphCPLMQA outperformed AlphaFold2 in terms of the MAE metric and demonstrated the ability to identify superior models compared to AlphaFold2. However, there are still challenges in effectively utilizing evaluation results to enhance model structure, and further advancements are needed in multimer model evaluation. We remain committed to making progress in these areas.

## Acknowledgments

This work was supported by the National Key R&D Program of China [2022ZD0115103], the National Nature Science Foundation of China [62173304, 62201506], the Key Project of Zhejiang Provincial Natural Science Foundation of China [LZ20F030002], the Basic Public Welfare Research Project of Zhejiang Province[LQ22F020028].

## Supplementary Information for

### Supplementary text

#### Supplementary Text S1 Equivariance proof process

In the network, feature *f* does not change its properties for orthogonal rotation matrix *R* and translation vector *T*. First, we define the model function as follows:

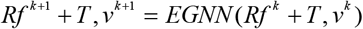

In our paper, the translation process of feature *f*:

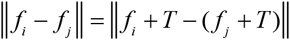

the rotation process of feature *f*:

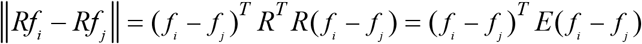

For the formula 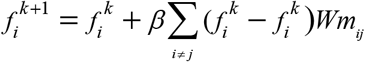 adds rotation and translation:

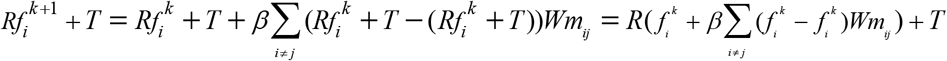

Therefore, with the rotation and translation of the input feature *f*, the same rotation and translation occur at the output end.

#### Supplementary Text S2 The multi-head attention

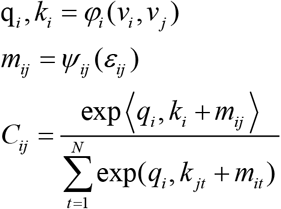

Where *φ*_*i*_, *ψ*_*ij*_ represent trainable linear functions, which map from *v*_*i*_, *v*_*j*_, *ε*_*ij*_ to q_*i*_, *k*_*i*_, *m*_*ij*_, respectively; 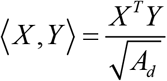 is used to scale the dot product attention operation between two matrices, where *A*_*d*_ is the dimension of attention.

#### Supplementary Text S3 The graph-equivariant update

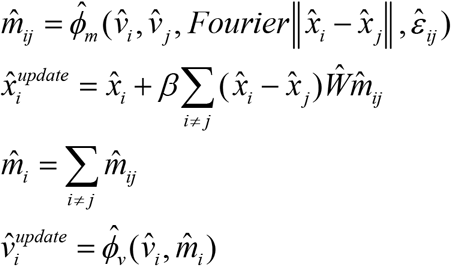

Where 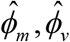 are graph network trainable linear layer; *Ŵ* is learnable matrix; *Fourier* 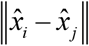 is fourier transform of the distance between node coordinates 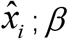 is (*N* − 1)^−1^.

#### Supplementary Text S4 Construction of local vectors and quadruples

we first construct a local coordinate system 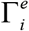 (the construction method is the same as *Γ*_*i*_). Secondly, the space vector *s*_*ij*_ between residue *i* and other residues *j* is calculated by rotating the transformation into the local coordinate system 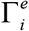 to obtain the local vector 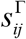. This indicates that the global spatial structure is mapped to each local space to enhance influence of protein local structure on overall topology. Moreover, in each local coordinate system 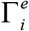, we rotate the coordinate system 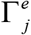 to get 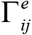 which is transformed into *Q*_ij_ by quadruple. The rotation transformation describes the relationship between each local spatial structure.

#### Supplementary Text S5 The process of updating the new nodes

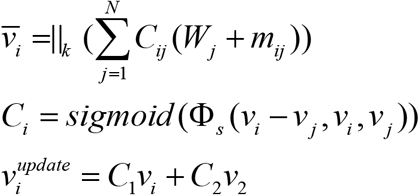

Where 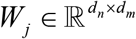 is attention head learning maxtrix; **Φ**_*s*_ is trainable linear function 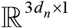. In the above, | denote splicing operation of multiple heads; *k* is number of heads; *C*_1_, *C*_2_ are *C*^*i*^ and 1− *C*^*i*^.

#### Supplementary Text S6 Local quality score

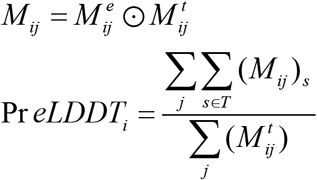

Where ⊙ denote distance error dot product distance thresholds to get the error within the threshold *s*(*T* ={0.5,1, 2, 4}).

#### Supplementary Text S7 Loss function

The total loss is the sum of the losses of the geometric constraint and quality assessment as follow:

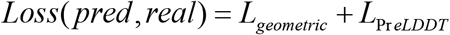

where *pred* and *real* are the predicted and real quality. Regarding *L*, it corresponds to the respective loss function.

#### Supplementary Text S8 Ablation studies for GraphCPLMQA

To investigate the impact of the components, we modified the full version of GraphCPLMQA by removing some features and changing the network architecture. Different network models were retrained to test the results and analyze the effect of these modifications. Firstly, GraphCPLMQA^1^ was created by replacing the transformer strategy inverted bottleneck of GraphCPLMQA with residual block in the decoding module, and GELU was replaced with ReLU to create GraphCPLMQA^1^. Regarding local metrics, there was a varying degree of decline in the prediction accuracy, while there was no significant change observed in global metrics. This indicated that the transformer strategy could further capture the local structural information. Secondly, the structural embeddings were removed from the language model in GraphCPLMQA^1^, leading to a decline in performance for GraphCPLMQA^2^. The high-dimensional structure features may imply some properties of protein structure that contribute to better learning of the network. Then, we changed the output mode of the encoding module (GraphCPLMQA^3^) and the connection architecture between modules (GraphCPLMQA^4^) on the GraphCPLMQA^2^ model. These operations may relatively weaken the sequence-structure relationship in the encoding module so that the encoding result has an impact on the decoding structure-quality relationship. Finally, based on GraphCPLMQA^2^model, the triangle position and residual-level contact order features were removed (GraphCPLMQA^5^). The reduction in accuracy on the local metrics implies that these characteristics can supplement the portrayal of the local structure.

## Supplementary figures

**Supplementary Figure S1.**
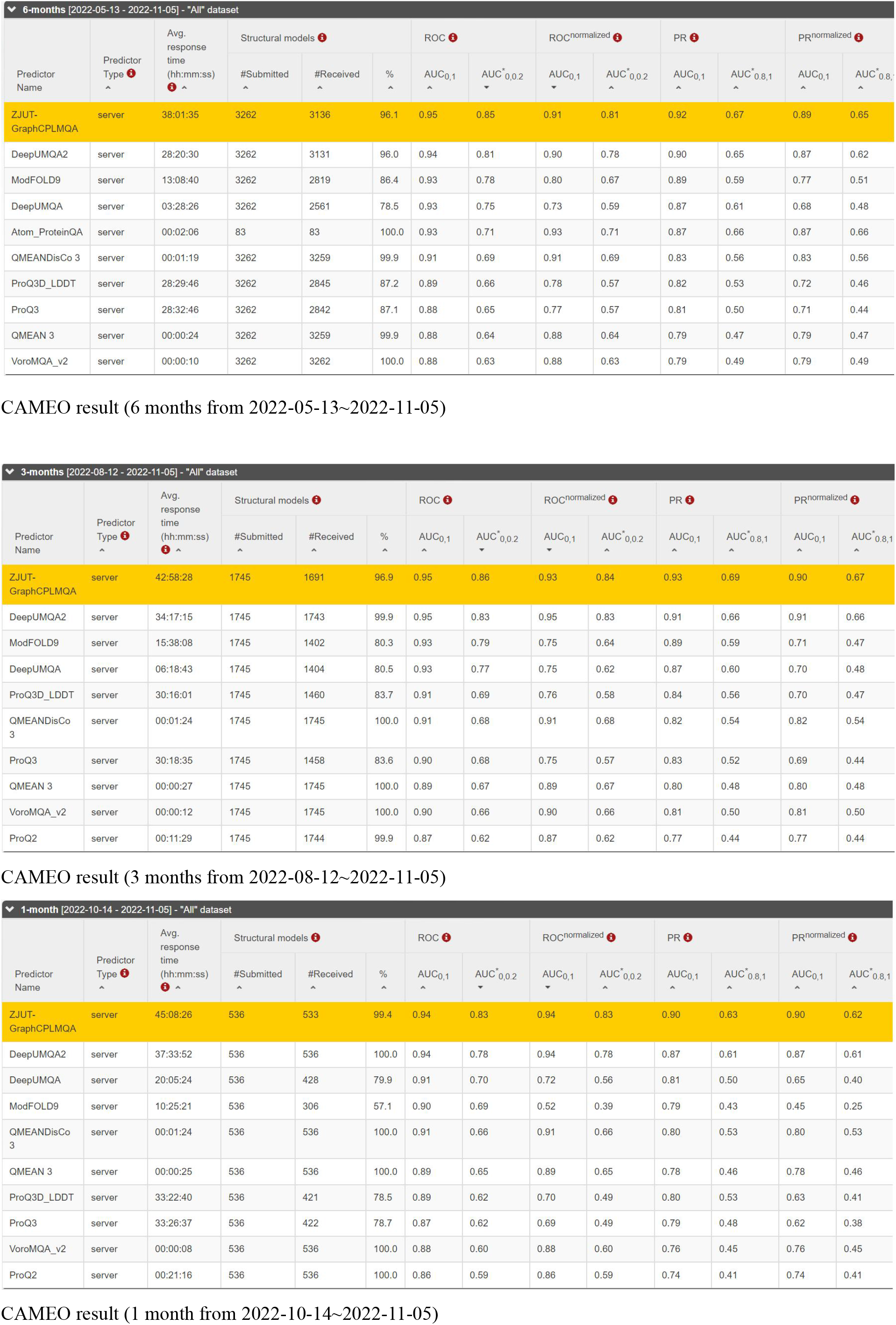
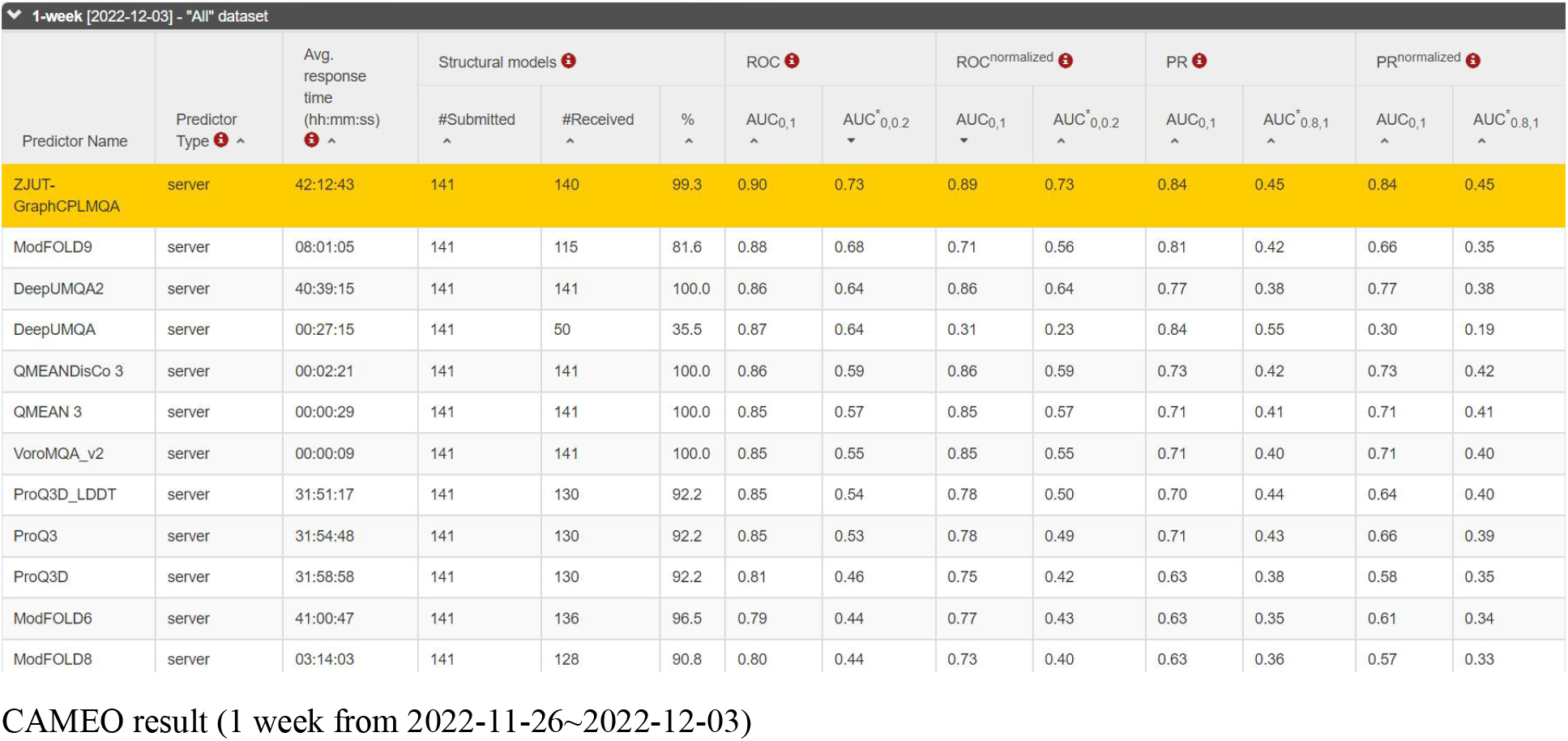

**Supplementary Figure S2.**
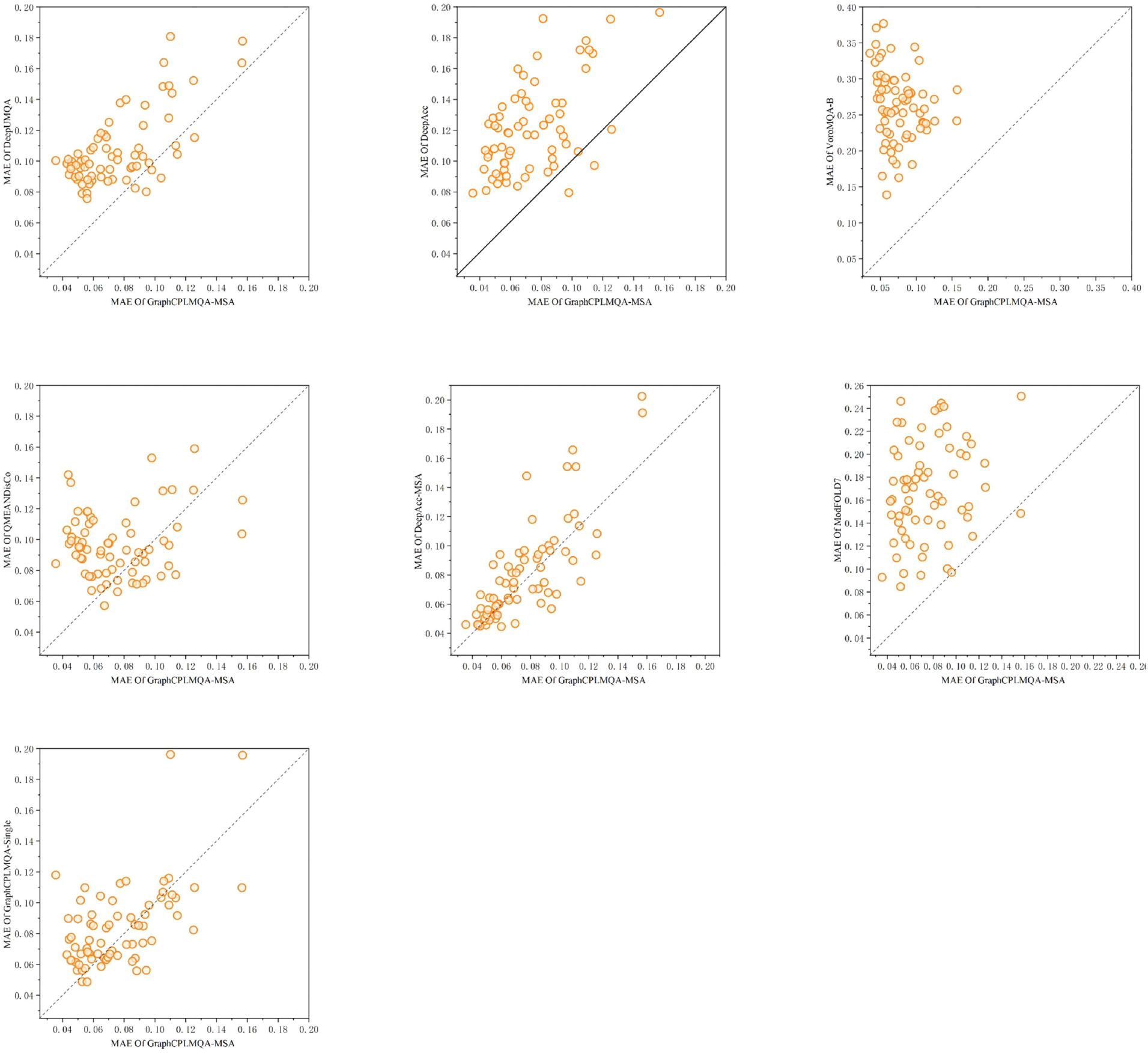
MAE of GraphCPLMQA (GraphCPLMQA-MSA) compared to other methods on CASP13

**Supplementary Figure S3.**
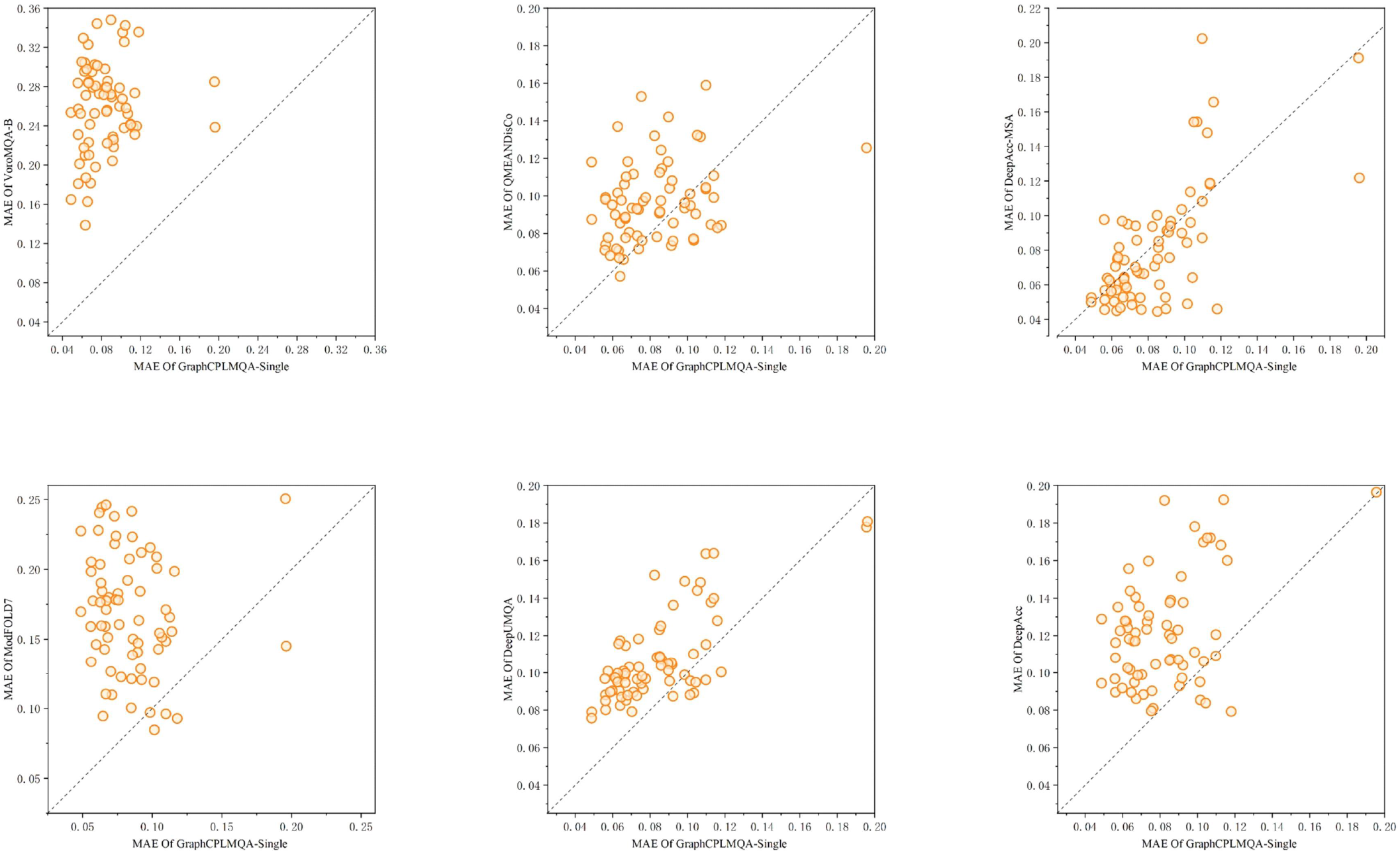
MAE of GraphCPLMQA-Single compared to other methods on CASP13.

**Supplementary Figure S4.**
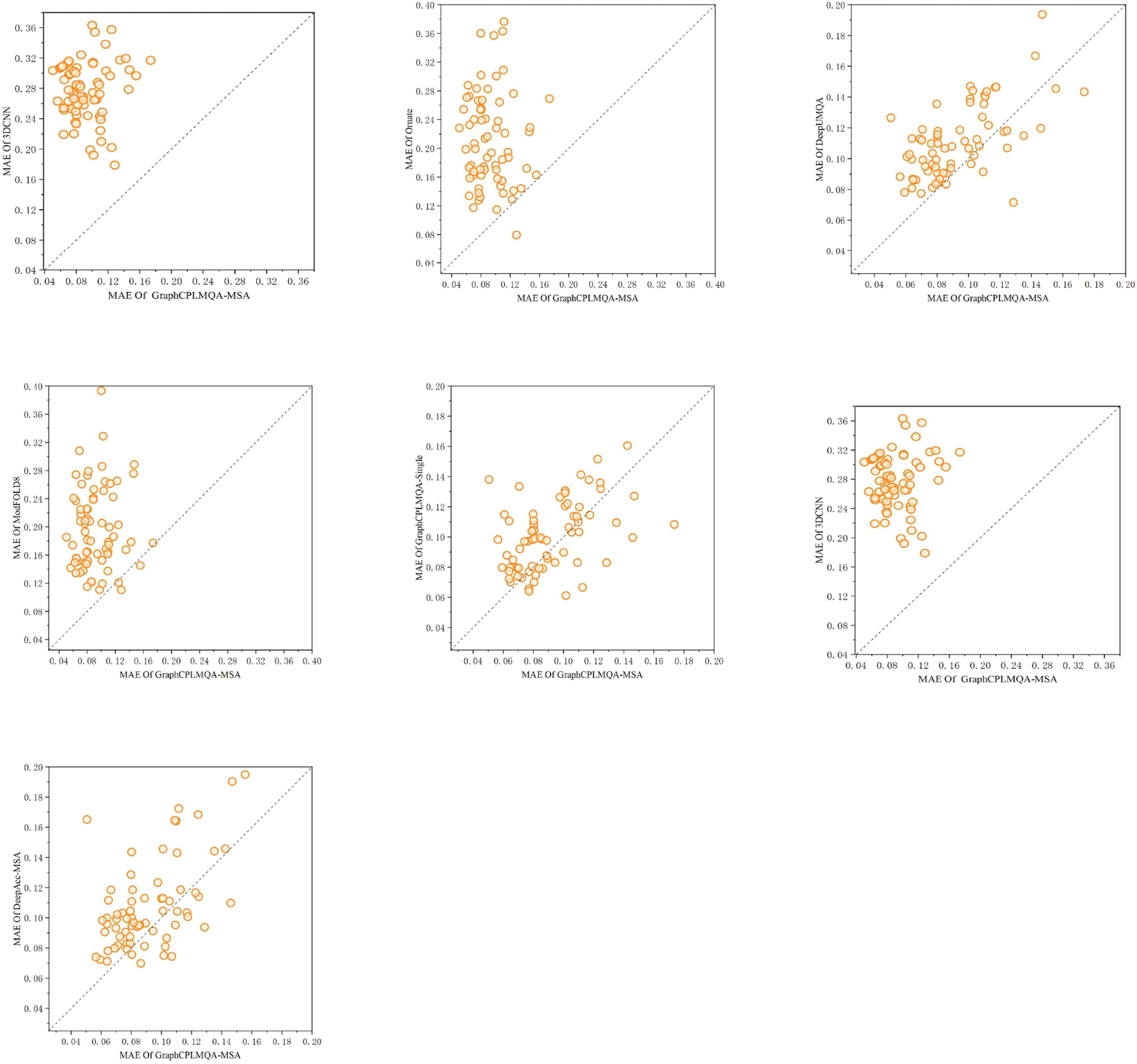
MAE of GraphCPLMQA (GraphCPLMQA-MSA) compared to other methods on CASP14

**Supplementary Figure S5.**
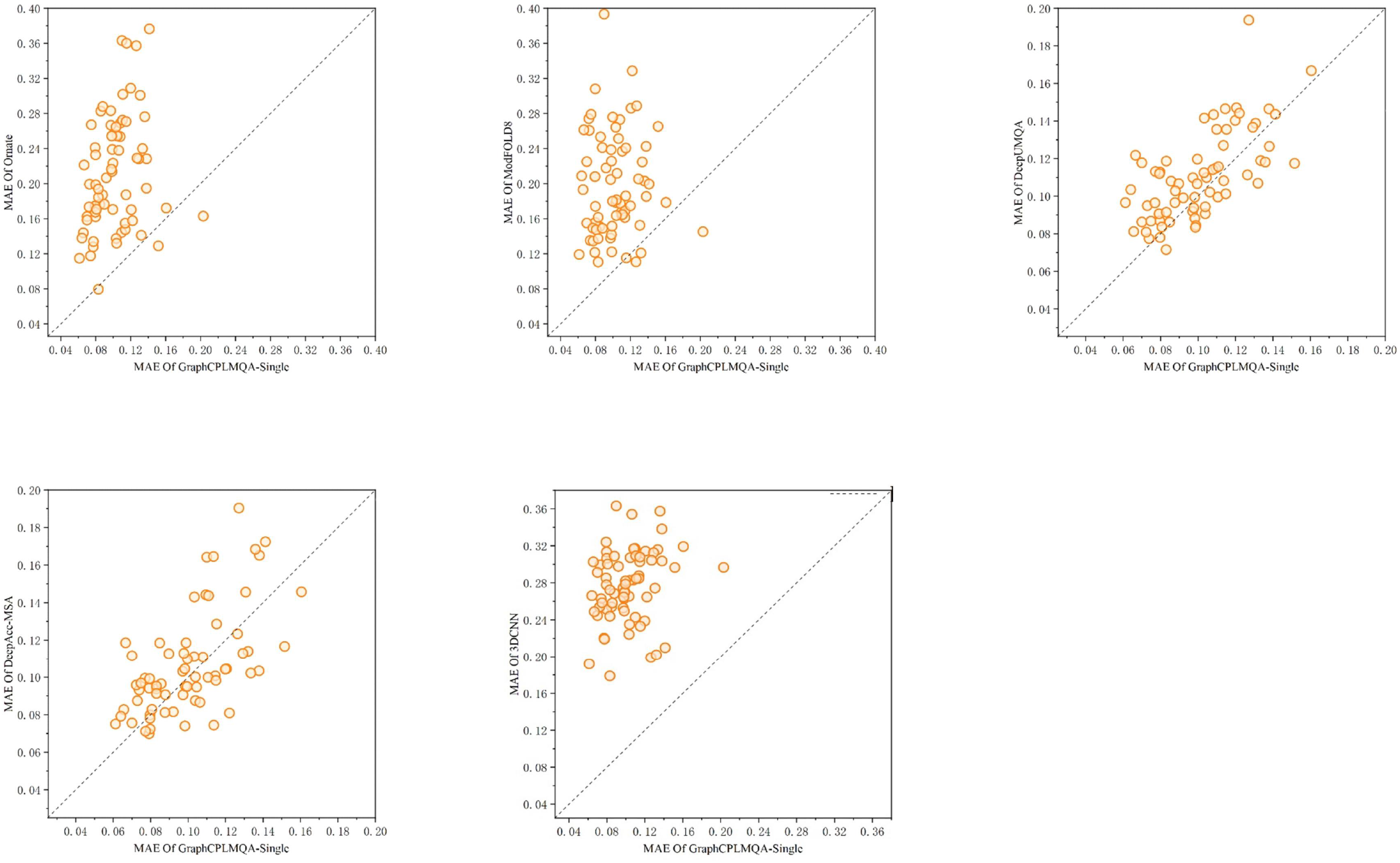
MAE of GraphCPLMQA-Single compared to other methods on CASP14

**Supplementary Figure S6.**
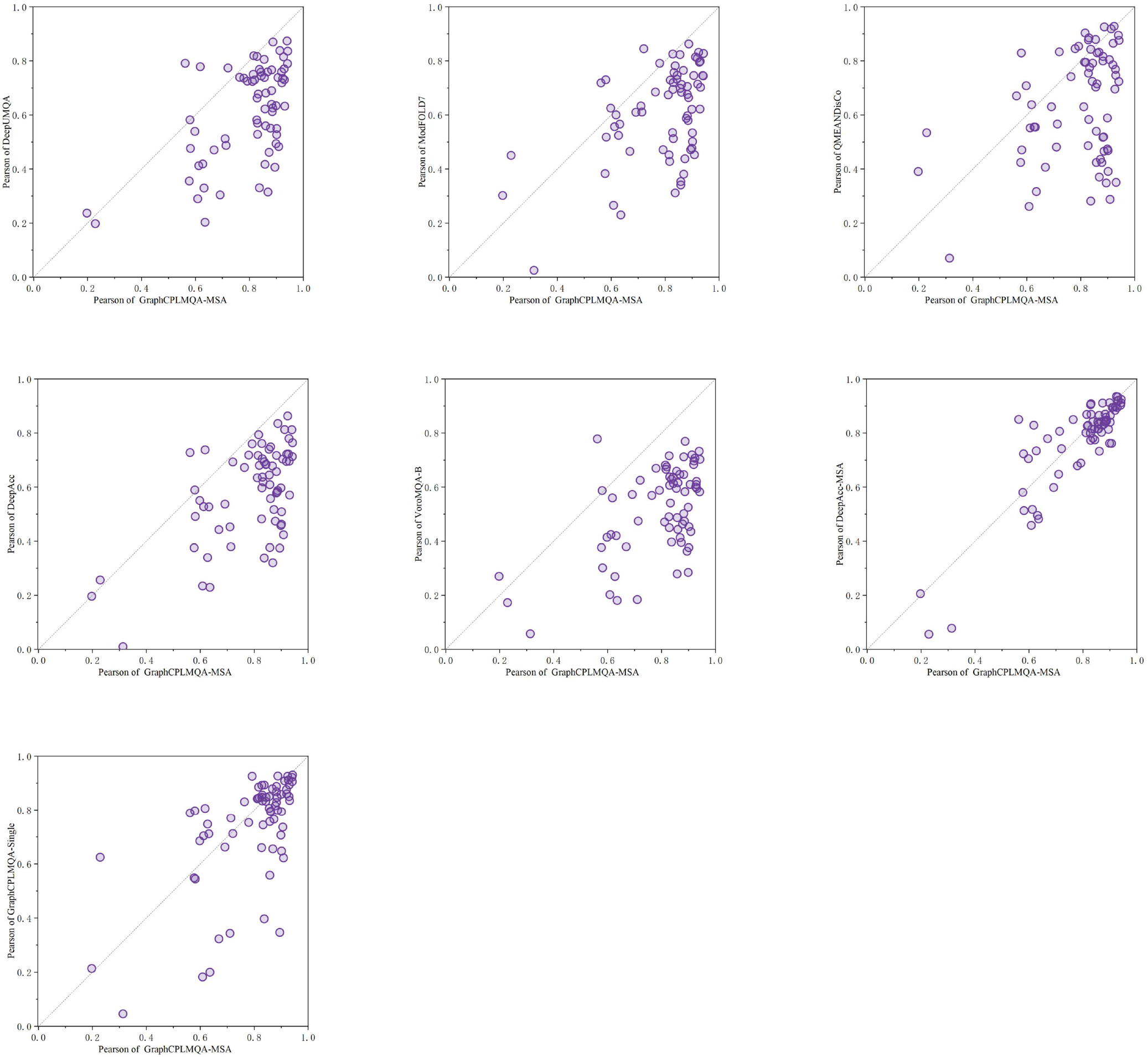
Pearson of GraphCPLMQA (GraphCPLMQA-MSA) compared to other methods on CASP13

**Supplementary Figure S7.**
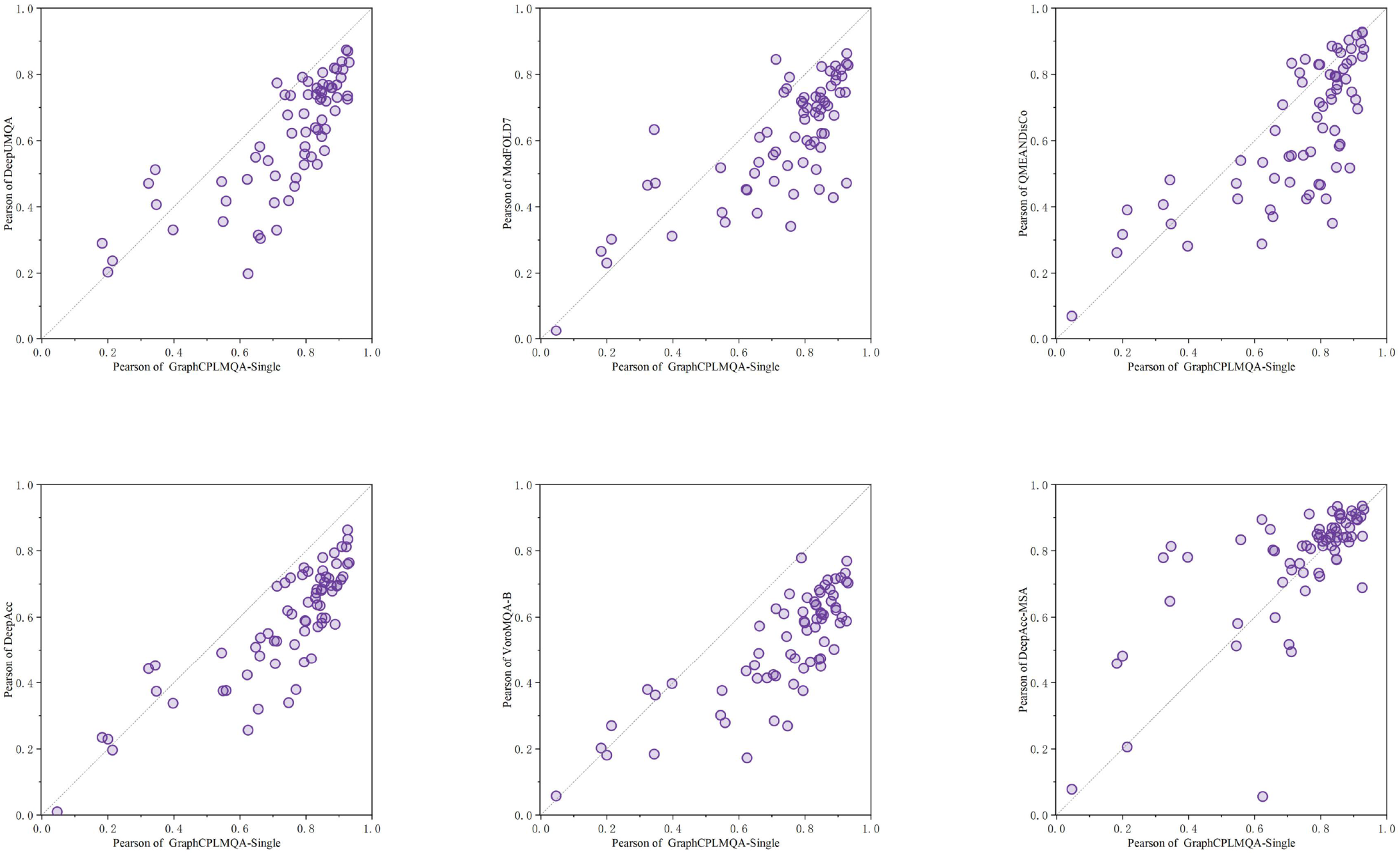
Pearson of GraphCPLMQA-Single compared to other methods on CASP13

**Supplementary Figure S7.**
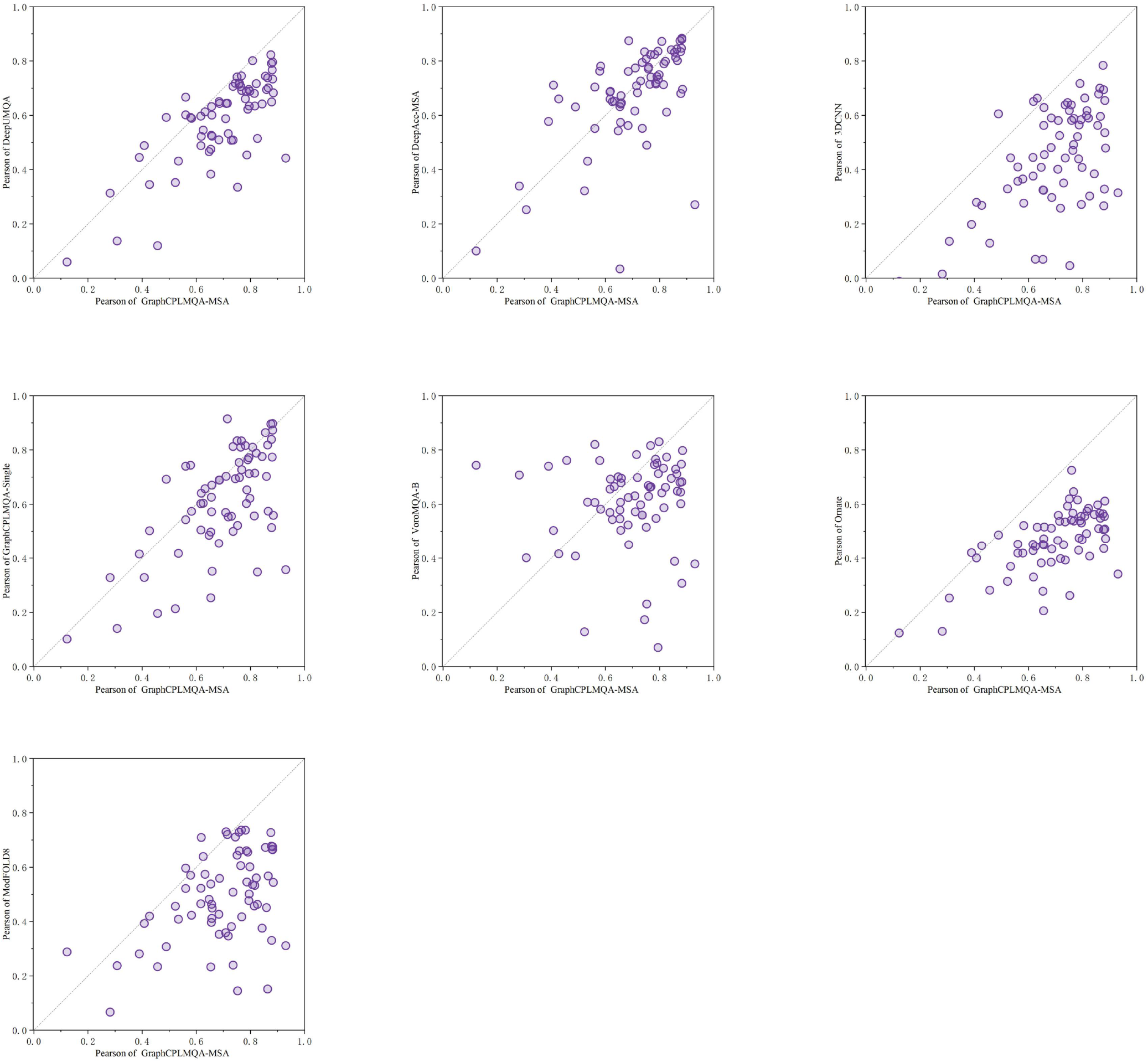
Pearson of GraphCPLMQA (GraphCPLMQA-MSA) compared to other methods on CASP14

**Supplementary Figure S8.**
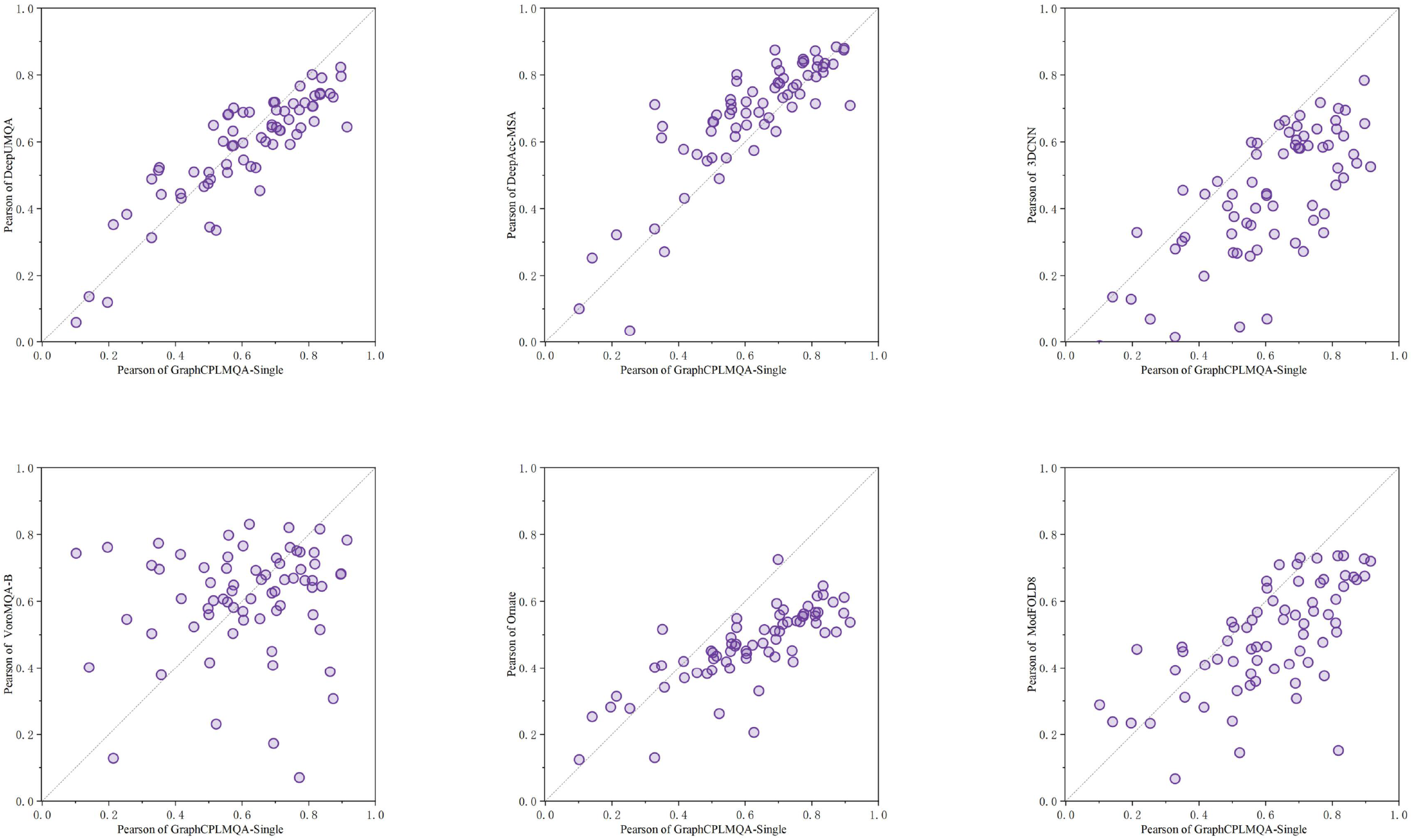
Pearson of GraphCPLMQA-Single compared to other methods on CASP14

**Supplementary Figure S9.**
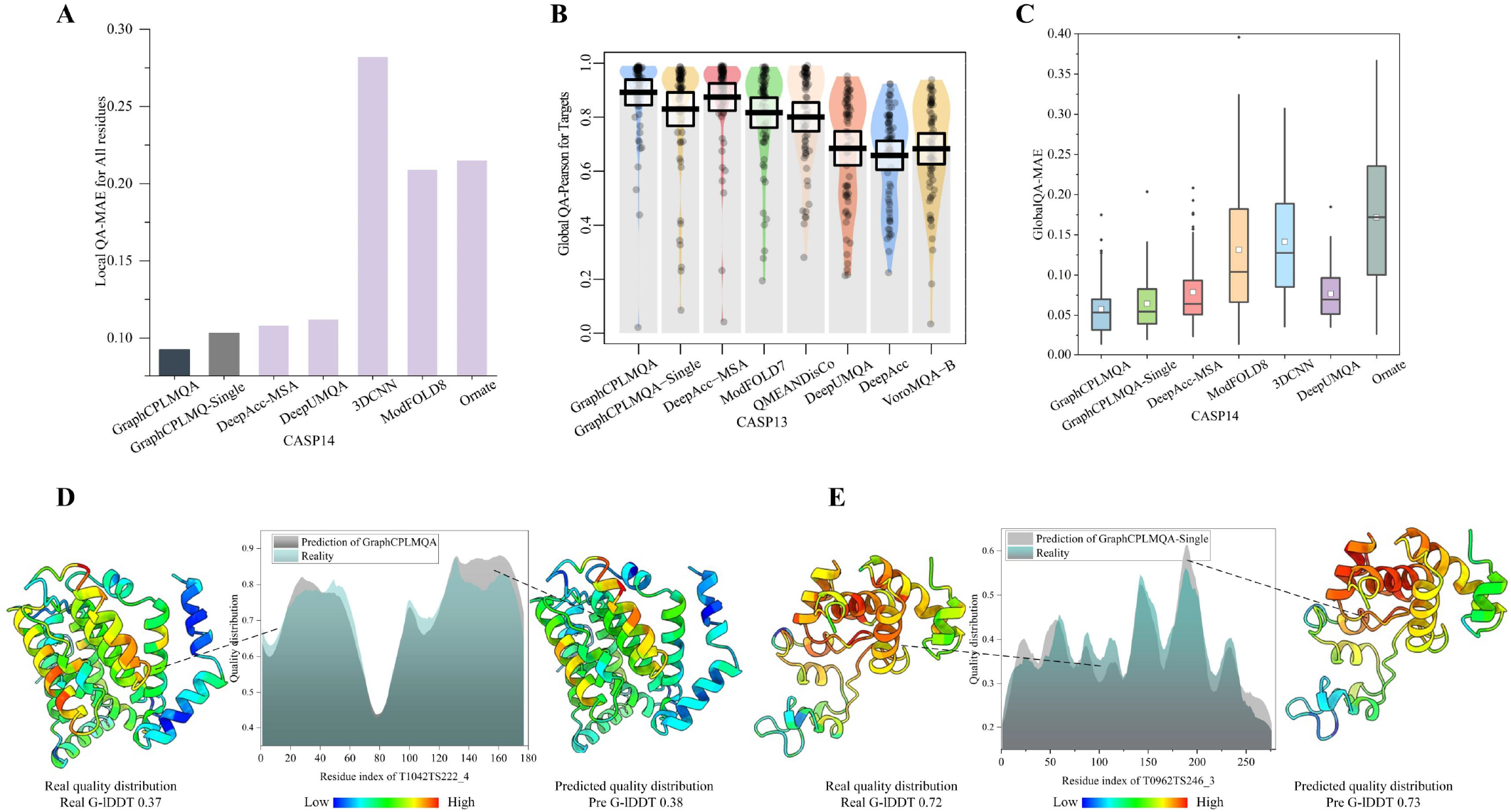
GraphCPLMQA compared to other methods on CASP monomer test set

**Supplementary Figure S10.**
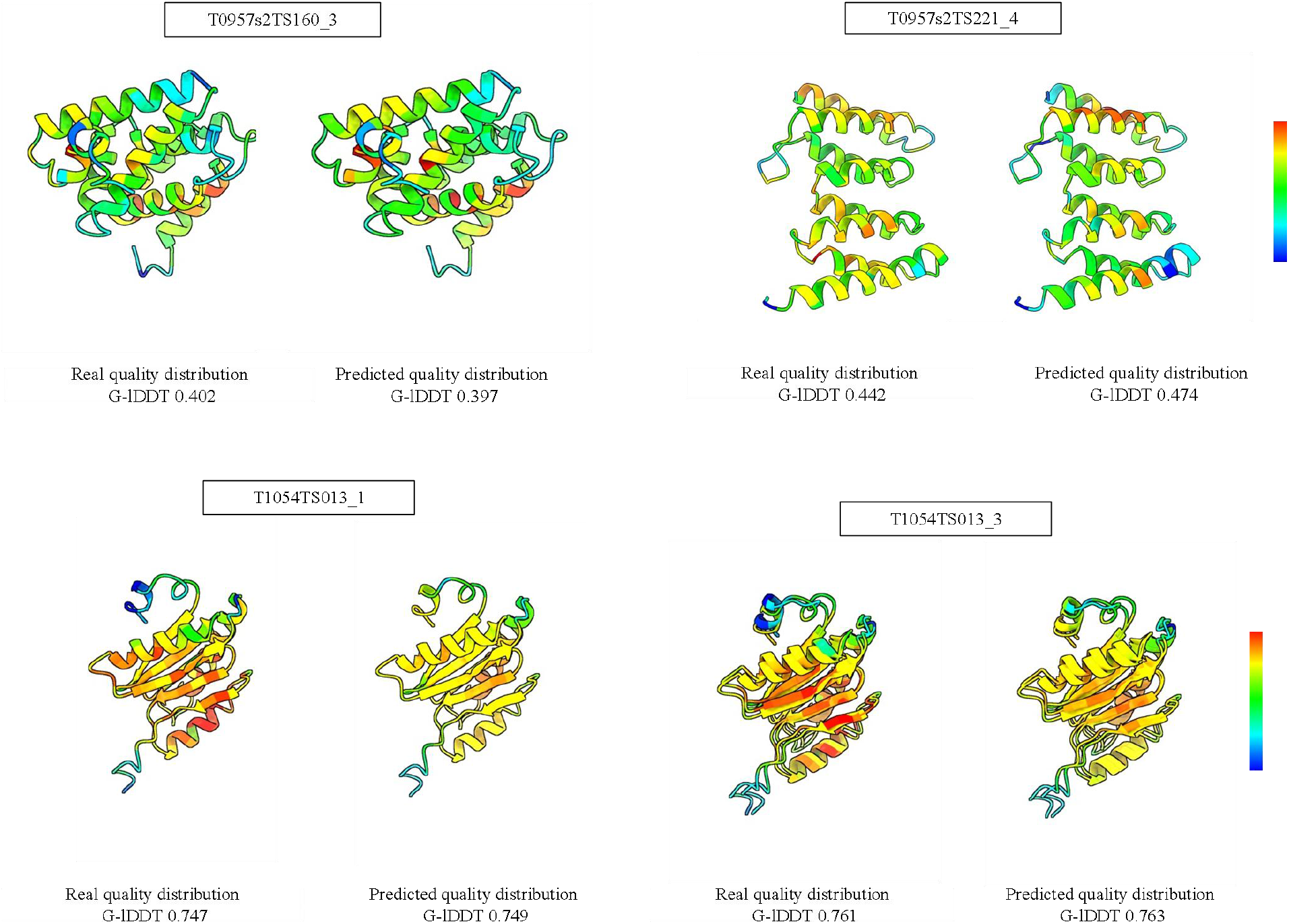
The prediction of GraphCPLMQA on the model of CASP dataset

**Supplementary Figure S11.**
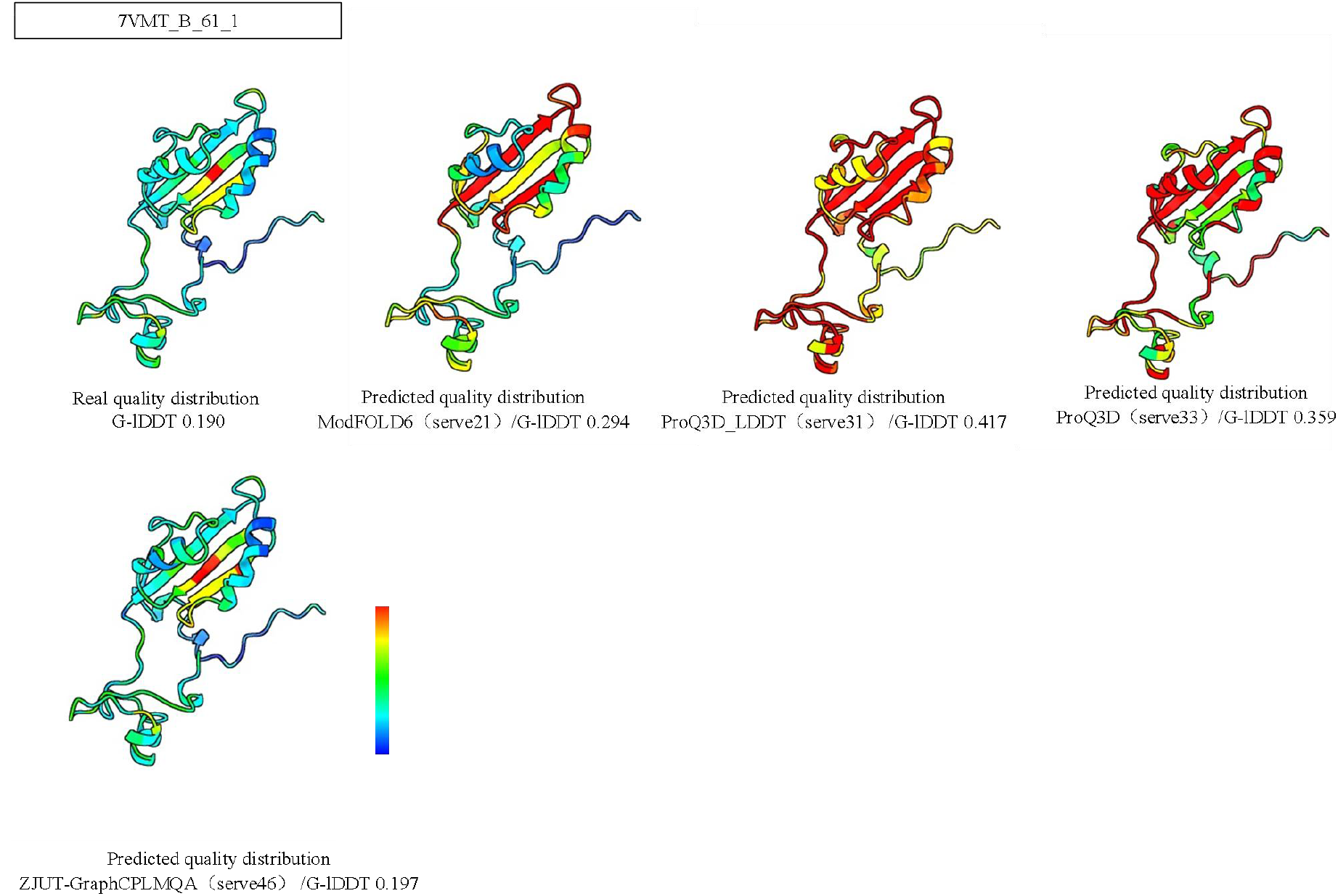
The prediction of ZJUT-GraphCPLMQA on the model of CAMEO dataset

**Supplementary Figure S12.**
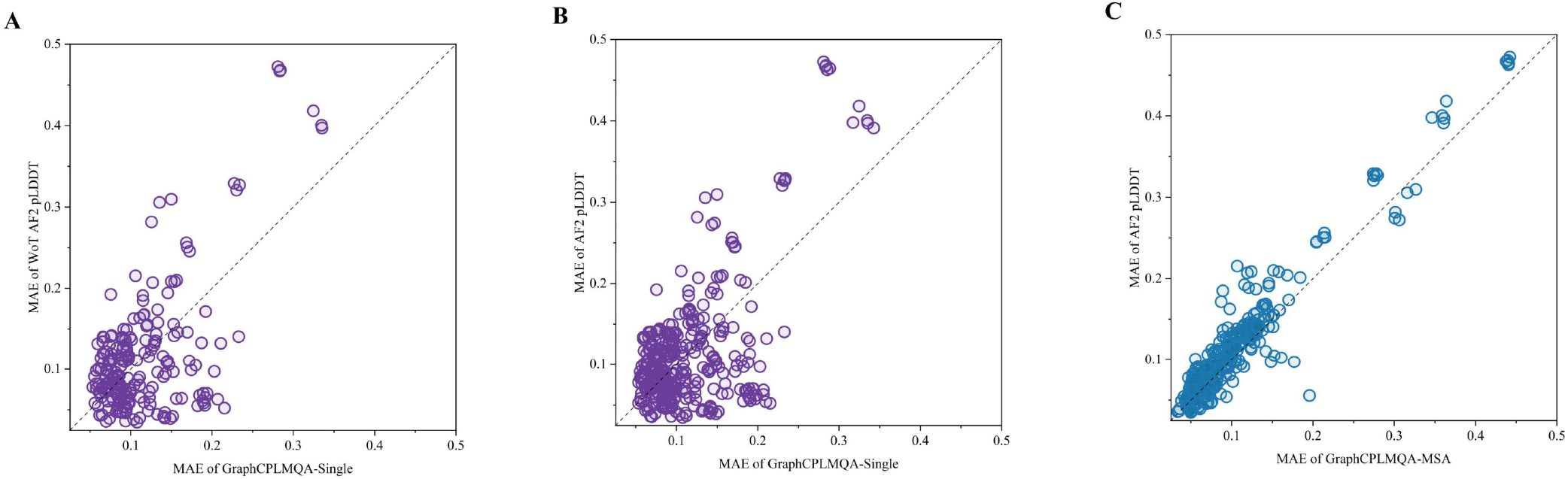
Results of GraphCPLMQA and GraphCPLMQA-Single on the AlphaFold Dataset

**Supplementary Figure S13.**
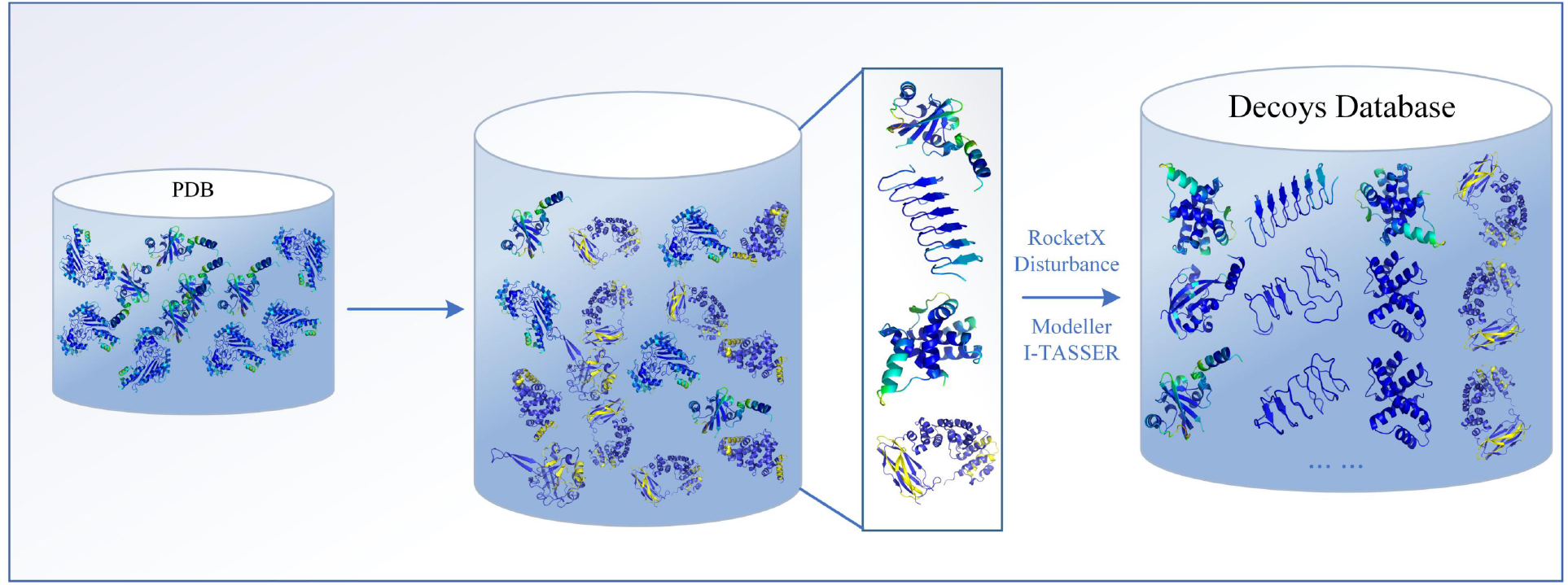
Schematic diagram of data set process construction

**Supplementary Figure S14.**
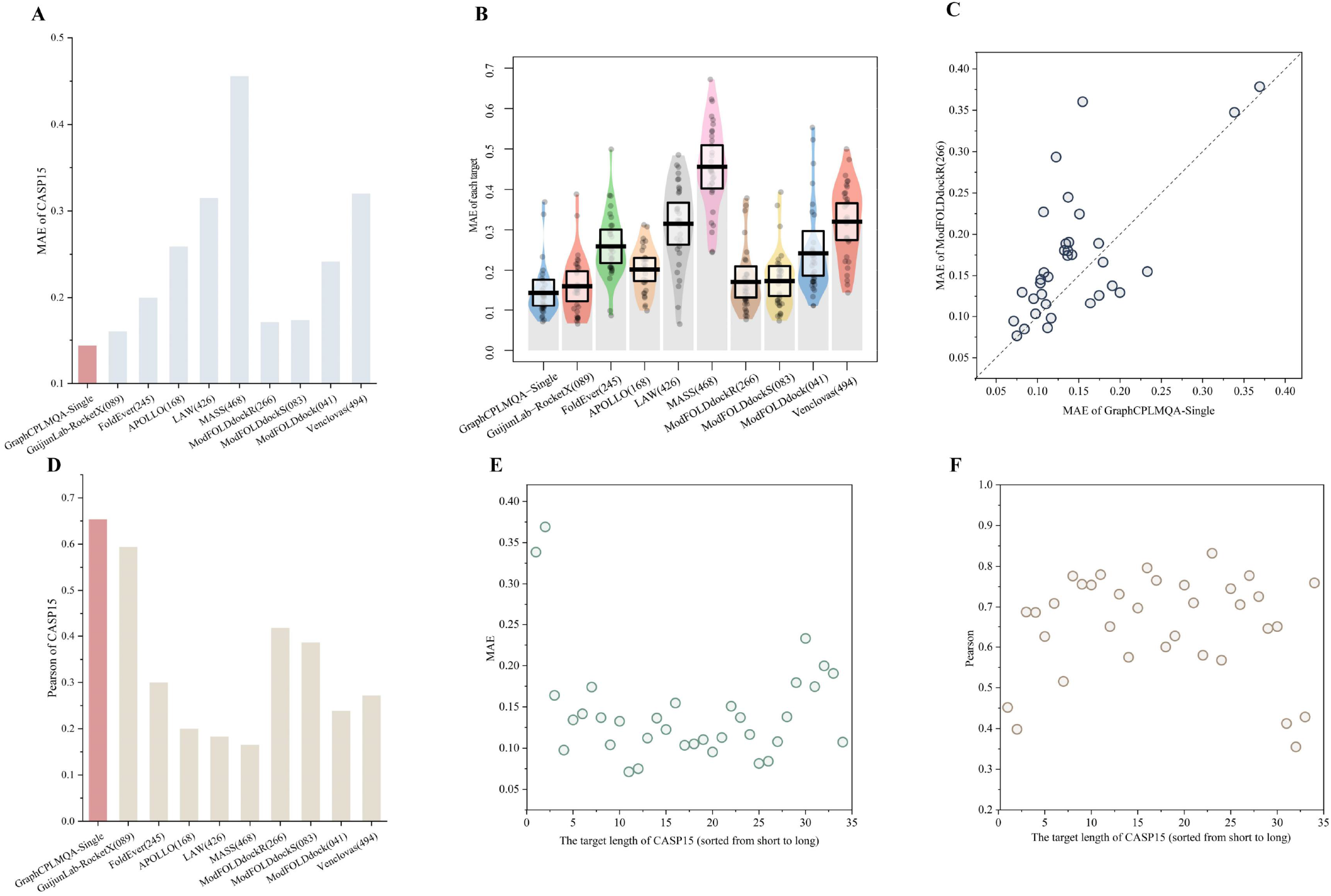
Results on the CASP complex test set

**Supplementary Figure S15.**
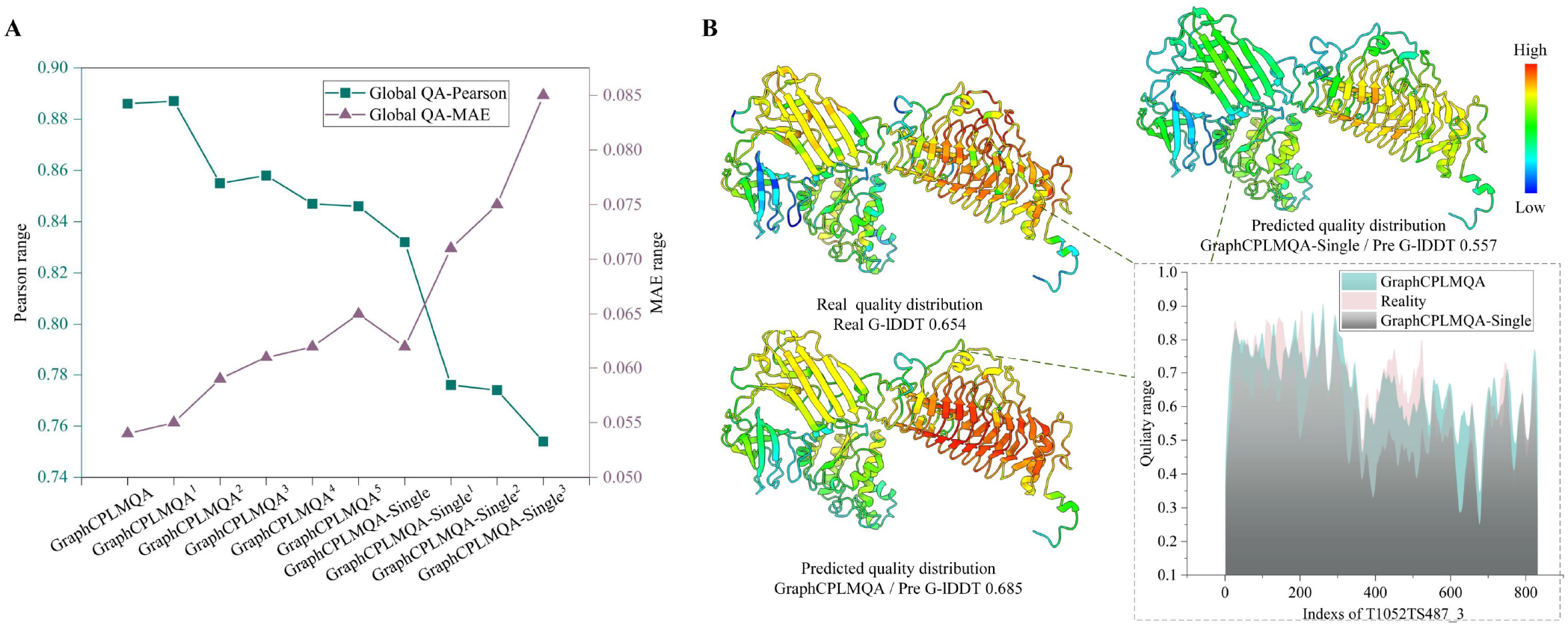
Ablation studies on CASP14

**Supplementary Figure S16.**
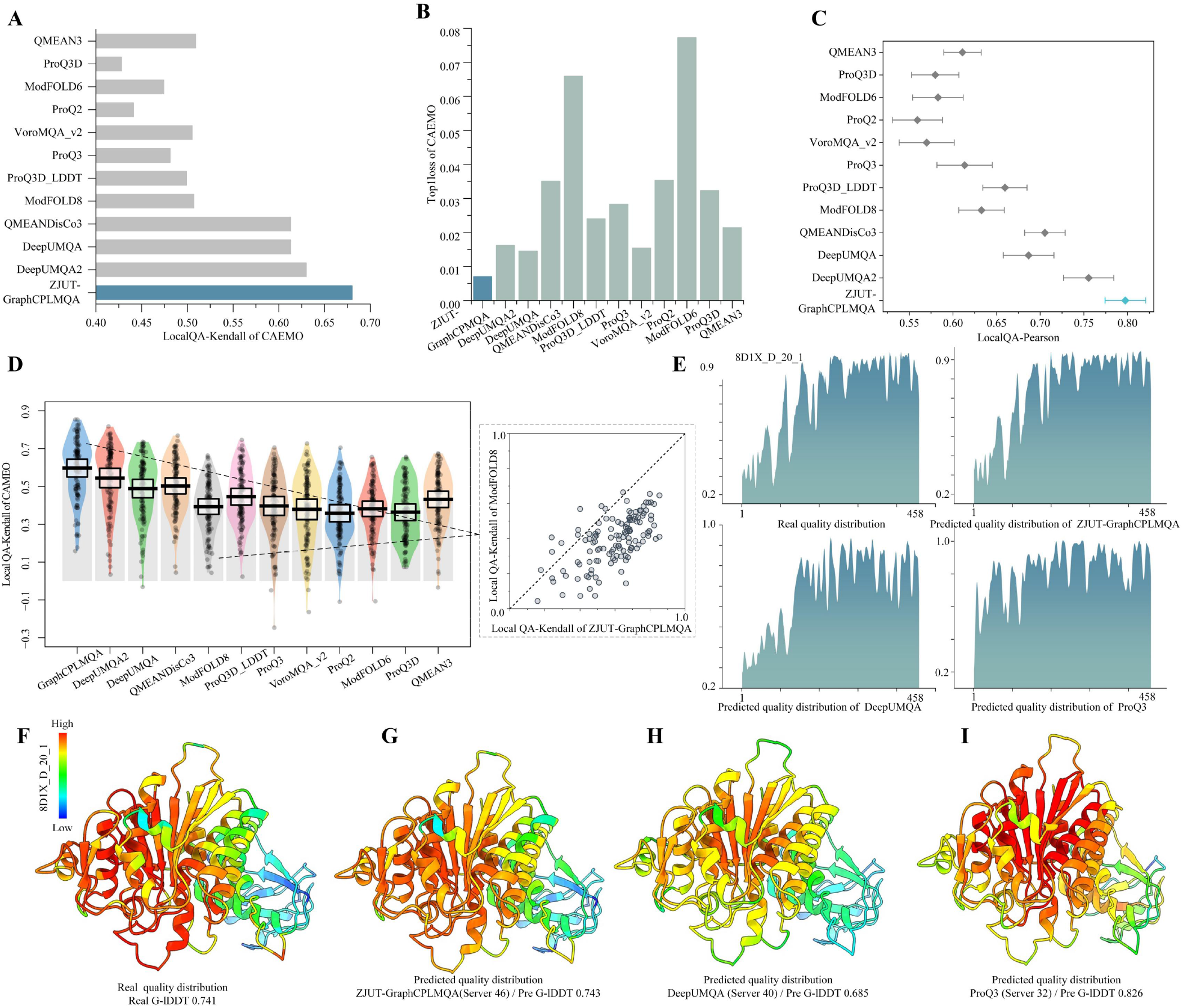
Result on CAEMO blind test

**Supplementary Figure S17.**
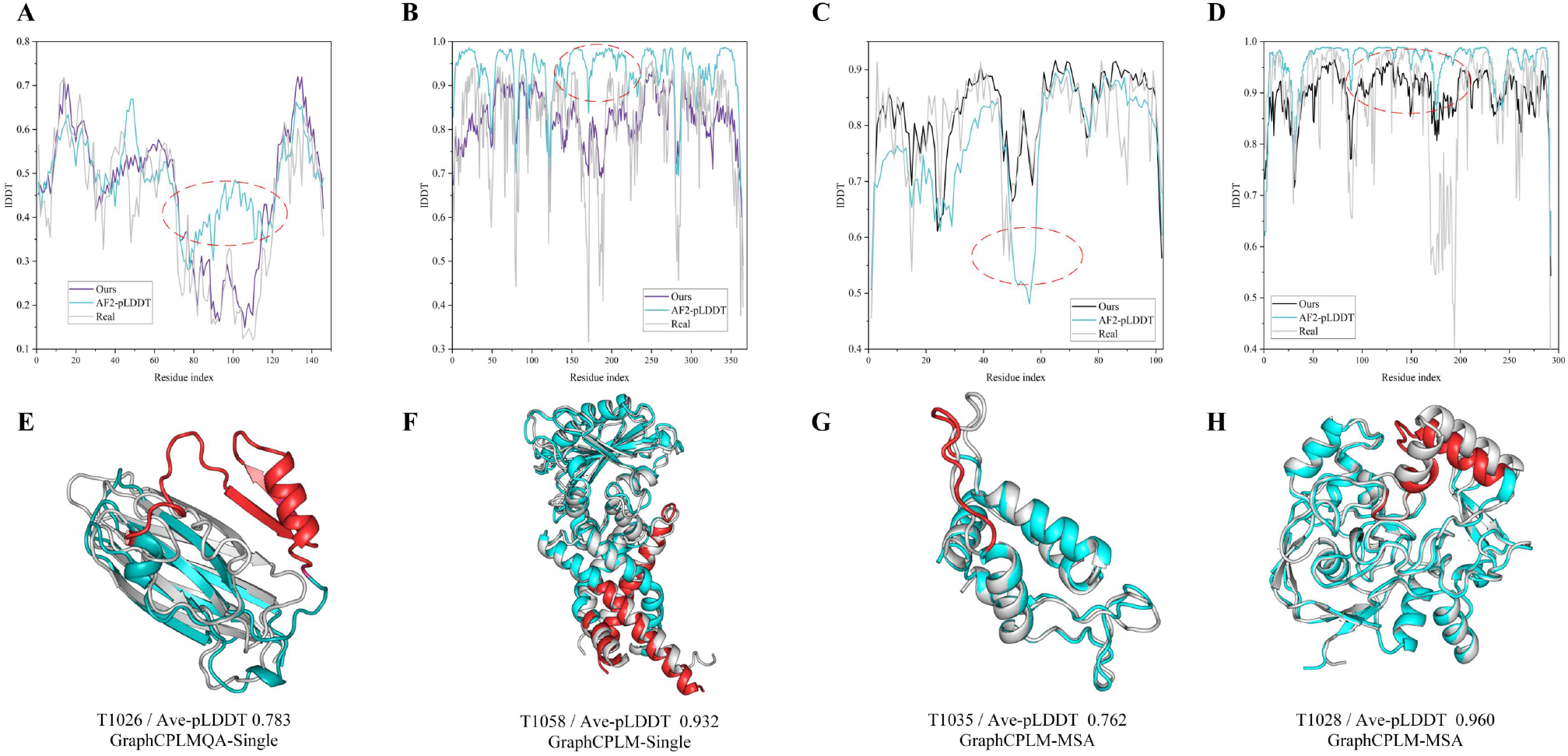
compare to AlphFold2. Quality analysis in the AlphaFold2 model. Figure (A∼D) corresponds to Figure (E∼H). In (E)- (H), gray represents the real structure, sky blue is the structure of AlphFold2, and red represents the region with relatively large folding error. For the medium and high-quality models of AlphaFold2, we analyzed the GraphCPLMQA prediction results and the pLDDT of AlphaFold2 (Supplementary Figure S17). On the AlphFold2 structure with medium quality (Supplementary Figure S17A, C), we could basically predict the distribution of local qulity and it was very close to the real distribution. The predicted local AlphaFold2 structure is not consistent with the native structure. In the case of AlphFold2 pLDDT, it is possible that AlphFold2 is not precise in local quality assessment, or even results in an opposite assessment, as indicated by the red area. This mets that the accuracy of AlphFold2 local structure prediction is closely related to the evaluation of local structure. To some extent, the pLDDT of AlphFold2 may not reflect the quality of the local structure. For the high-quality AlphFold2 structure (Supplementary Figure S17B, D), our evaluation results were more consistent with the distribution of the real quality. However, most predicted results of AlphFold2 were higher than the real quality.

## Supplementary tables

**Supplementary Table S1.** 70 targets protein of CASP13

|  |  |  |  |  |  |  |
| --- | --- | --- | --- | --- | --- | --- |
| T0991 | T0981 | T0957s2 | T0988 | T1017s1 | T0958 | T0957s1 |
| T0984 | T1019s2 | T1004 | T0970 | T0977 | T0978 | T0974s2 |
| T0967 | T0955 | T0961 | T0998 | T1010 | T0968s1 | T1021s2 |
| T0974s1 | T0971 | T1000 | T0953s2 | T0979 | T0987 | T0954 |
| T0983 | T0986s2 | T0951 | T0963 | T0966 | T1020 | T1015s1 |
| T0982 | T1008 | T0989 | T1014 | T1016 | T0985 | T0990 |
| T0976 | T1018 | T1015s2 | T0949 | T1019s1 | T0973 | T0962 |
| T0980s1 | T0992 | T0950 | T0993s1 | T0986s1 | T1013 | T1002 |
| T1011 | T0969 | T0964 | T1017s2 | T0997 | T0980s2 | T0960 |
| T1009 | T0953s1 | T0975 | T1021s1 | T0965 | T0968s2 | T0959 |

**Supplementary Table S2.**
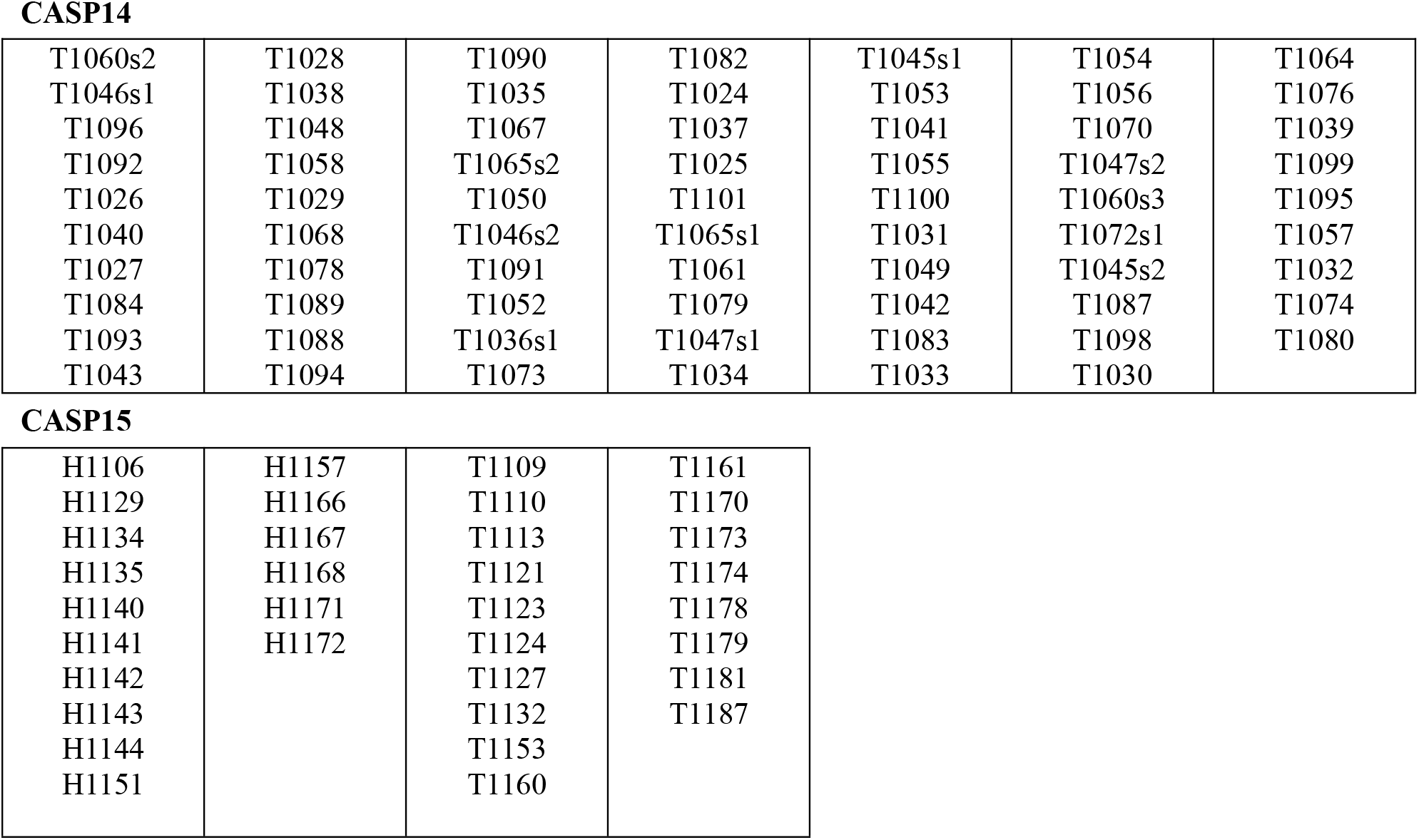
69 and 34 targets protein of CASP14 and CASP15

CASP14
|  |  |  |  |  |  |  |
| --- | --- | --- | --- | --- | --- | --- |
| T1060s2 | T1028 | T1090 | T1082 | T1045s1 | T1054 | T1064 |
| T1046s1 | T1038 | T1035 | T1024 | T1053 | T1056 | T1076 |
| T1096 | T1048 | T1067 | T1037 | T1041 | T1070 | T1039 |
| T1092 | T1058 | T1065s2 | T1025 | T1055 | T1047s2 | T1099 |
| T1026 | T1029 | T1050 | T1101 | T1100 | T1060s3 | T1095 |
| T1040 | T1068 | T1046s2 | T1065s1 | T1031 | T1072s1 | T1057 |
| T1027 | T1078 | T1091 | T1061 | T1049 | T1045s2 | T1032 |
| T1084 | T1089 | T1052 | T1079 | T1042 | T1087 | T1074 |
| T1093 | T1088 | T1036s1 | T1047s1 | T1083 | T1098 | T1080 |
| T1043 | T1094 | T1073 | T1034 | T1033 | T1030 |  |

CASP15
|  |  |  |  |
| --- | --- | --- | --- |
| H1106 | H1157 | T1109 | T1161 |
| H1129 | H1166 | T1110 | T1170 |
| H1134 | H1167 | T1113 | T1173 |
| H1135 | H1168 | T1121 | T1174 |
| H1140 | H1171 | T1123 | T1178 |
| H1141 | H1172 | T1124 | T1179 |
| H1142 |  | T1127 | T1181 |
| H1143 |  | T1132 | T1187 |
| H1144 |  | T1153 |  |
| H1151 |  | T1160 |  |

**Supplementary Table S3.**
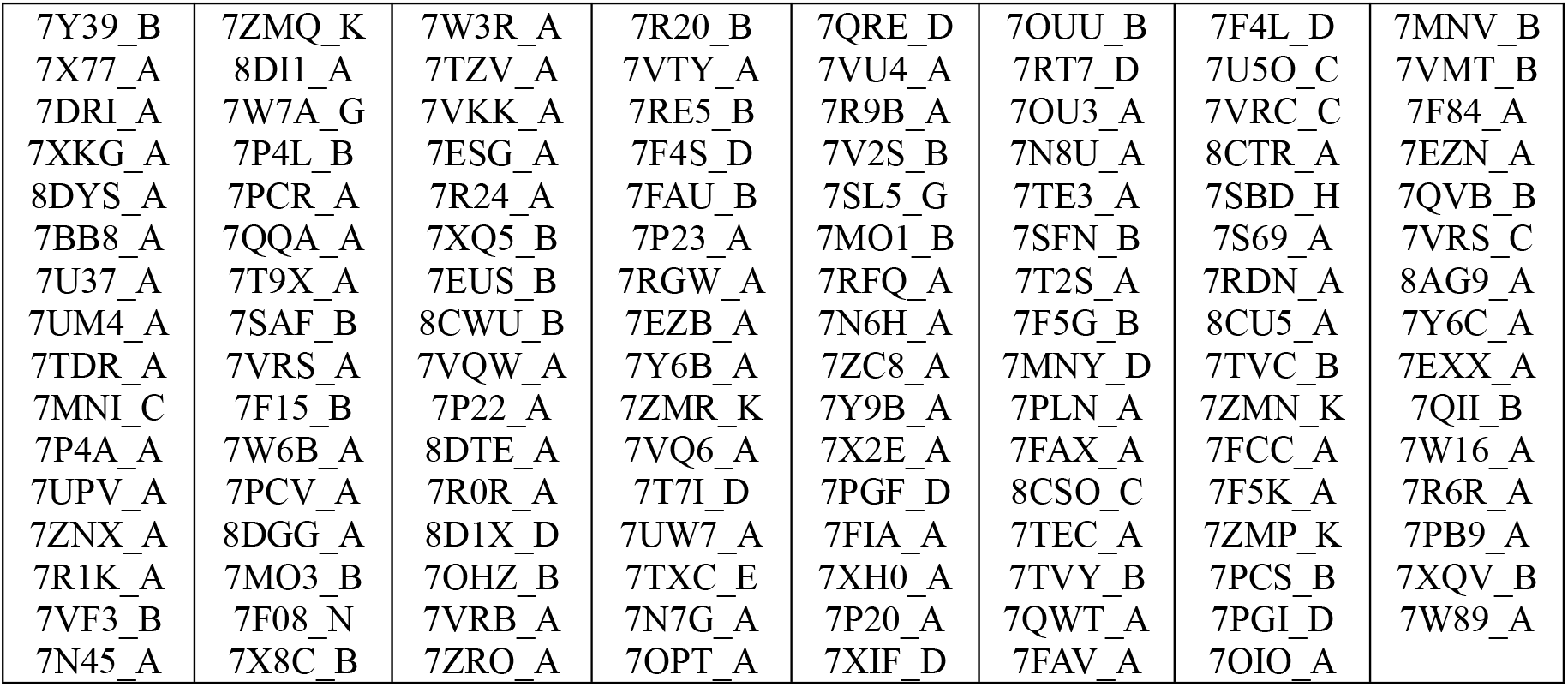
127 targets protein of CAEMO (5.20∼8.13)

**Supplementary Table S4.** ESM language model

| Type | model | input | Out(dim) | Layer | Dataset |
| --- | --- | --- | --- | --- | --- |
| ESM-2 | esm2_t33_650M_UR50D | single sequence | 1280 | 33 | UR50 |
| ESM-MSA-1b[1] | esm_msa1b_t12_100M_UR50S | MSA | 768,144 (attention) | 12 | UR50 |
| ESM-IF1[2] | esm_if1_gvp4_t16_142M_UR50 | backbone atomic coordinates | 512 | 20 | CATH + UR50 |
| ESM-1v | esm1v_t33_650M_UR90S_1 | single sequence | 1280 | 33 | UR90 |
| ESM-1b | esm1b_t33_650M_UR50S | single sequence | 1280 | 33 | UR50 |

**Note:** UR is UniRef database, CATH is a classification of protein structures downloaded from the Protein Data Bank. The ESM-IF1 inverse folding model is built for predicting protein sequences from their backbone atom coordinates.

**Supplementary Table S5.**
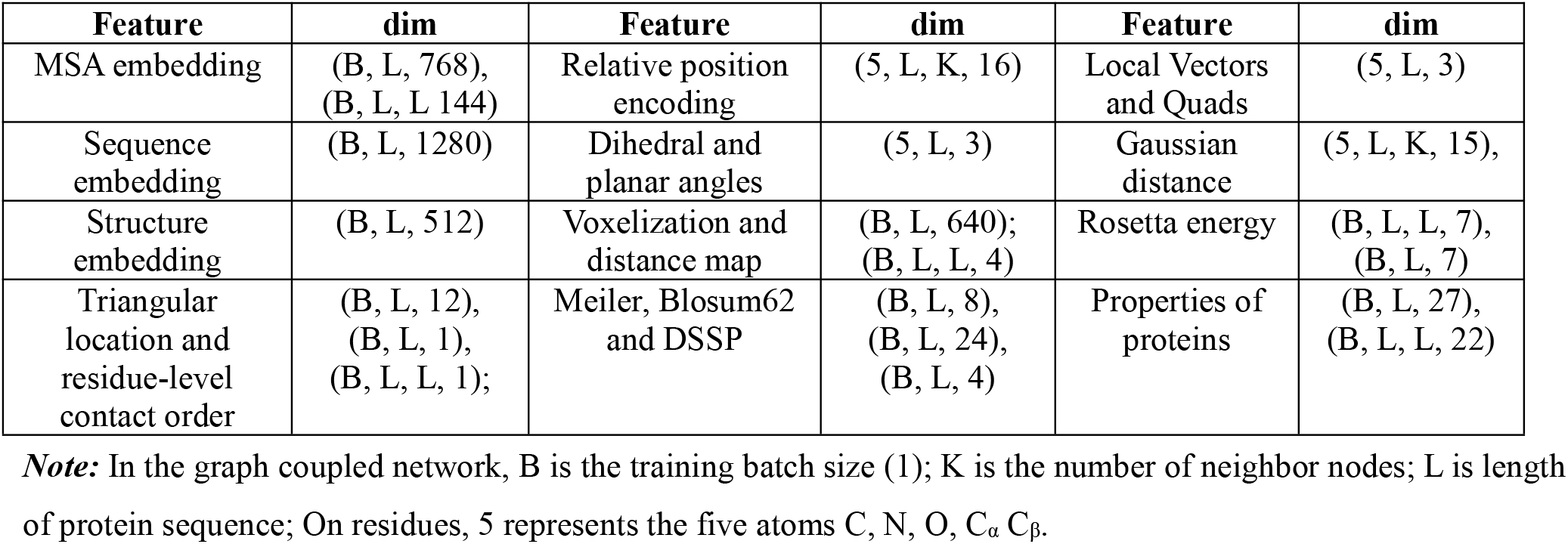
Feature of GraphCPLMQA

**Supplementary Table S6.** Parameters of Graph Coupled Network

| Parameters | Description | Value |
| --- | --- | --- |
| GT-$d_n$ | Graph Transformer node dimension | 64 |
| GT-$d_e$ | Graph Transformer edge dimension | 64 |
| GT-$k$ | Graph Transformer attention head number | 4 |
| GT-$d_{head}$ | Graph Transformer attention head dimension | 32 |
| GT-$d_{depth}$ | Graph Transformer layer number | 4 |
| Tri-in-$d_n$ | In-Triangular multiplication node dimension | 64 |
| Tri-in-$d_e$ | In-Triangular multiplication edge dimension | 64 |
| Tri-in-$d_{hid}$ | In-Triangular multiplication hidden dimension | 128 |
| Tri-out-$d_n$ | Out-Triangular multiplication node dimension | 64 |
| Tri-out-$d_e$ | Out-Triangular multiplication edge dimension | 64 |
| Tri-out-$d_{hid}$ | Out-Triangular multiplication hidden dimension | 128 |
| Seq-block | Sequence embedding block | 3 |
| IPA- $d_n$ | IPA node dimension | 128 |
| IPA- $d_{depth}$ | IPA layer number | 4 |
| IPA- $k$ | IPA head number | 8 |
| IPA-block | IPA block | 3 |
| EGNN-globalattention- $d_n$ | EGNN global attention | 64 |
| EGNN-$d_n$ | EGNN node dimension | 64 |
| ConvT-er | Transformer-based convolution dilation rate | 1,2,4,8,16 |
| ConvT-IB | Inverted bottleneck | 128-256-128 |
| ConvT-$d_e$ | Transformer-based convolution dimension | 128 |
| ConvT-$s$ | Transformer-based convolution kernel size | 1-3-1 |
| ConvT-Main-block | Transformer-based main convolution block | 3 |
| ConvT-Error-block | Transformer-based error convolution block | 3 |
| ConvT-Threshold-block | Transformer-based threshold convolution block | 3 |

**Supplementary Table S7.** Comparison of GraphCPLMQA with other methods on 9390 models of CASP13

| Methods | Local QA |  |  |  |  | Global QA |  |  |  |  |
| --- | --- | --- | --- | --- | --- | --- | --- | --- | --- | --- |
|  | Person | Kendall | AUC | MSE | MAE | Person | Kendall | AUC | MAE | Top1loss |
| **GraphCPLMQA** | **0.860** | **0.681** | **0.930** | **0.012** | **0.079** | **0.927** | **0.777** | **0.962** | **0.044** | **0.028** |
| **GraphCPLMQA-Single** | **0.837** | **0.648** | **0.932** | **0.014** | **0.087** | **0.893** | **0.734** | **0.949** | **0.050** | **0.047** |
| DeepAcc-MSA | 0.831 | 0.645 | 0.914 | 0.016 | 0.091 | 0.902 | 0.743 | 0.949 | 0.052 | 0.033 |
| QMEANDisCo | 0.813 | 0.609 | 0.926 | 0.017 | 0.096 | 0.897 | 0.710 | 0.958 | 0.057 | 0.061 |
| DeepUMQA | 0.748 | 0.546 | 0.899 | 0.022 | 0.112 | 0.830 | 0.621 | 0.933 | 0.074 | 0.065 |
| ModFOLD7 | 0.736 | 0.552 | 0.882 | 0.045 | 0.168 | 0.826 | 0.653 | 0.893 | 0.123 | 0.081 |
| DeepAcc | 0.735 | 0.533 | 0.886 | 0.030 | 0.135 | 0.797 | 0.589 | 0.909 | 0.103 | 0.058 |
| VoroMQA-B | 0.577 | 0.424 | 0.810 | 0.101 | 0.266 | 0.643 | 0.507 | 0.847 | 0.217 | 0.070 |

**Note:** Results obtained by calculating all residues for all models (in the same way as the official CAMEO calculation). Pearson and Kendall represent the trend between predicted quality and true quality. AUC (Area under curve) is the area enclosed by the coordinate axis under the ROC curve. MSE and MAE reflect the difference between the predicted value and the true value. Top1loss is an indicator for selecting a model, reflecting the error between the predicted best structure and the real best structure in candidate models.

**Supplementary Table S8.** Comparison of GraphCPLMQA with other methods on 9645 models of CASP14

| Methods | Local QA |  |  |  |  | Global QA |  |  |  |  |
| --- | --- | --- | --- | --- | --- | --- | --- | --- | --- | --- |
|  | Person | Kendall | AUC | MSE | MAE | Person | Kendall | AUC | MAE | Top1loss |
| **GraphCPLMA** | **0.784** | **0.594** | **0.882** | **0.015** | **0.092** | **0.886** | **0.721** | **0.948** | **0.054** | **0.031** |
| DeepAcc-MSA | 0.756 | 0.571 | 0.876 | 0.020 | 0.108 | 0.858 | 0.691 | 0.941 | 0.078 | 0.039 |
| **GraphCPLMQA-Single** | **0.733** | **0.544** | **0.865** | **0.018** | **0.103** | **0.832** | **0.654** | **0.939** | **0.062** | **0.048** |
| DeepUMQA | 0.701 | 0.511 | 0.851 | 0.021 | 0.112 | 0.795 | 0.600 | 0.917 | 0.073 | 0.042 |
| 3DCNN | 0.583 | 0.422 | 0.798 | 0.106 | 0.282 | 0.791 | 0.599 | 0.915 | 0.141 | 0.058 |
| ModFOLD8 | 0.561 | 0.396 | 0.779 | 0.064 | 0.209 | 0.622 | 0.428 | 0.830 | 0.126 | 0.075 |
| Ornate | 0.474 | 0.335 | 0.742 | 0.068 | 0.215 | 0.665 | 0.497 | 0.856 | 0.173 | 0.056 |

**Note:** Results obtained by calculating all residues for all models (in the same way as the official CAMEO calculation).

**Supplementary Table S9.** GraphCPLMQA ablation experiments on 9390 models of CASP13

| Methods | Local QA |  |  |  |  | Global QA |  |  |  |  |
| --- | --- | --- | --- | --- | --- | --- | --- | --- | --- | --- |
|  | Person | Kendall | AUC | MSE | MAE | Person | Kendall | AUC | MAE | Top1loss |
| GraphCPLMA | 0.860 | 0.681 | 0.930 | 0.012 | 0.079 | 0.927 | 0.777 | 0.962 | 0.044 | 0.028 |
| GraphCPLMA<sup>1</sup> | 0.831 | 0.648 | 0.919 | 0.014 | 0.087 | 0.906 | 0.748 | 0.951 | 0.049 | 0.047 |
| GraphCPLMA<sup>2</sup> | 0.823 | 0.633 | 0.924 | 0.015 | 0.088 | 0.908 | 0.748 | 0.964 | 0.049 | 0.048 |
| GraphCPLMA<sup>3</sup> | 0.817 | 0.629 | 0.915 | 0.015 | 0.093 | 0.900 | 0.740 | 0.959 | 0.054 | 0.050 |
| GraphCPLMA<sup>4</sup> | 0.808 | 0.626 | 0.910 | 0.016 | 0.092 | 0.899 | 0.736 | 0.963 | 0.053 | 0.043 |
| GraphCPLMA<sup>5</sup> | 0.807 | 0.616 | 0.905 | 0.016 | 0.091 | 0.899 | 0.736 | 0.953 | 0.048 | 0.047 |

**Supplementary Table S10.** GraphCPLMQA-Single ablation experiments on 9390 models of CASP13

| Methods | Local QA |  |  |  |  | Global QA |  |  |  |  |
| --- | --- | --- | --- | --- | --- | --- | --- | --- | --- | --- |
|  | Person | Kendall | AUC | MSE | MAE | Person | Kendall | AUC | MAE | Top1loss |
| GraphCPLMA-Single | 0.837 | 0.648 | 0.932 | 0.014 | 0.087 | 0.893 | 0.734 | 0.947 | 0.050 | 0.047 |
| GraphCPLMA-Single<sup>1</sup> | 0.796 | 0.601 | 0.908 | 0.017 | 0.099 | 0.857 | 0.675 | 0.929 | 0.061 | 0.056 |
| GraphCPLMA-Single<sup>2</sup> | 0.791 | 0.592 | 0.912 | 0.017 | 0.101 | 0.853 | 0.661 | 0.946 | 0.064 | 0.055 |
| GraphCPLMA-Single<sup>3</sup> | 0.747 | 0.551 | 0.879 | 0.025 | 0.118 | 0.817 | 0.620 | 0.911 | 0.079 | 0.053 |

**Supplementary Table S11.** GraphCPLMQA ablation experiments on 9645 models of CASP14

| Methods | Local QA |  |  |  |  | Global QA |  |  |  |  |
| --- | --- | --- | --- | --- | --- | --- | --- | --- | --- | --- |
|  | Person | Kendall | AUC | MSE | MAE | Person | Kendall | AUC | MAE | Top1loss |
| GraphCPLMA | 0.784 | 0.594 | 0.882 | 0.015 | 0.092 | 0.886 | 0.721 | 0.948 | 0.054 | 0.031 |
| GraphCPLMA<sup>1</sup> | 0.772 | 0.585 | 0.880 | 0.016 | 0.096 | 0.887 | 0.722 | 0.947 | 0.055 | 0.025 |
| GraphCPLMA<sup>2</sup> | 0.750 | 0.562 | 0.865 | 0.017 | 0.098 | 0.855 | 0.683 | 0.928 | 0.059 | 0.035 |
| GraphCPLMA<sup>3</sup> | 0.748 | 0.558 | 0.867 | 0.017 | 0.099 | 0.858 | 0.692 | 0.929 | 0.061 | 0.025 |
| GraphCPLMA<sup>4</sup> | 0.726 | 0.539 | 0.855 | 0.020 | 0.105 | 0.847 | 0.681 | 0.933 | 0.062 | 0.043 |
| GraphCPLMA<sup>5</sup> | 0.725 | 0.536 | 0.852 | 0.019 | 0.106 | 0.846 | 0.666 | 0.919 | 0.065 | 0.042 |

**Supplementary Table S12.** GraphCPLMQA-Single ablation experiments on 9645 models of CASP14

| Methods | Local QA |  |  |  |  | Global QA |  |  |  |  |
| --- | --- | --- | --- | --- | --- | --- | --- | --- | --- | --- |
|  | Person | Kendall | AUC | MSE | MAE | Person | Kendall | AUC | MAE | Top1loss |
| GraphCPLMA-Single | 0.733 | 0.544 | 0.865 | 0.018 | 0.103 | 0.832 | 0.654 | 0.939 | 0.062 | 0.048 |
| GraphCPLMA-Single<sup>1</sup> | 0.693 | 0.506 | 0.845 | 0.020 | 0.110 | 0.776 | 0.586 | 0.906 | 0.071 | 0.048 |
| GraphCPLMA-Single<sup>2</sup> | 0.675 | 0.492 | 0.835 | 0.023 | 0.114 | 0.774 | 0.591 | 0.902 | 0.075 | 0.056 |
| GraphCPLMA-Single<sup>3</sup> | 0.656 | 0.474 | 0.823 | 0.028 | 0.130 | 0.754 | 0.567 | 0.903 | 0.085 | 0.050 |

**Supplementary Table S13.** Results of ZJUT-GraphCPLMQA (server 46) on CAMEO blind test set

| Methods | Local QA |  |  |  |  | Global QA |  |  |  |  |
| --- | --- | --- | --- | --- | --- | --- | --- | --- | --- | --- |
|  | Person | Kendall | AUC | MSE | MAE | Person | Kendall | AUC | MAE | Top1loss |
| **ZJUT-GraphCPLMA** | **0.891** | **0.680** | **0.942** | **0.015** | **0.081** | **0.924** | **0.741** | **0.967** | **0.077** | **0.007** |
| DeepUMQA2 | 0.878 | 0.630 | 0.941 | 0.013 | 0.083 | 0.908 | 0.687 | 0.966 | 0.066 | 0.016 |
| DeepUMQA | 0.835 | 0.613 | 0.923 | 0.017 | 0.099 | 0.874 | 0.684 | 0.956 | 0.074 | 0.015 |
| QMEANDisCo3 | 0.829 | 0.613 | 0.925 | / | / | 0.885 | 0.664 | 0.962 | 0.354 | 0.035 |
| ModFOLD8 | 0.771 | 0.507 | 0.894 | 0.026 | 0.127 | 0.824 | 0.557 | 0.920 | 0.094 | 0.066 |
| ProQ3D_LDDT | 0.769 | 0.499 | 0.887 | 0.038 | 0.151 | 0.824 | 0.530 | 0.904 | 0.137 | 0.024 |
| ProQ3 | 0.749 | 0.481 | 0.881 | 0.039 | 0.156 | 0.825 | 0.559 | 0.914 | 0.129 | 0.028 |
| VoroMQA_v2 | 0.733 | 0.505 | 0.880 | 0.051 | 0.176 | 0.824 | 0.584 | 0.926 | 0.166 | 0.015 |
| ProQ2 | 0.717 | 0.441 | 0.859 | 0.050 | 0.176 | 0.812 | 0.520 | 0.900 | 0.159 | 0.035 |
| ModFOLD6 | 0.716 | 0.474 | 0.877 | 0.052 | 0.182 | 0.811 | 0.536 | 0.906 | 0.160 | 0.077 |
| ProQ3D | 0.708 | 0.428 | 0.853 | 0.037 | 0.151 | 0.811 | 0.509 | 0.892 | 0.112 | 0.032 |
| QMEAN3 | 0.705 | 0.509 | 0.886 | 0.165 | 0.333 | 0.807 | 0.611 | 0.921 | 0.357 | 0.021 |
| VoroMQ_sw5 | 0.596 | 0.395 | 0.828 | 0.051 | 0.188 | 0.766 | 0.515 | 0.878 | 0.160 | 0.023 |

**Supplementary Table S14.** Performance of the method on homologous and heterologous complexes

| Methods | Homo-Pearson | Heter-Pearson | Homo-MAE | Heter-MAE |
| --- | --- | --- | --- | --- |
| **GraphCPLMA-single** | **0.655** | **0.652** | **0.144** | **0.143** |
| GuijunLab-RocketX | 0.600 | 0.586 | 0.169 | 0.150 |
| APOLLO | 0.242 | 0.147 | 0.251 | 0.269 |
| FoldEver | 0.354 | 0.247 | 0.185 | 0.219 |
| LAW | 0.100 | 0.289 | 0.309 | 0.323 |
| MASS | 0.142 | 0.196 | 0.438 | 0.479 |
| ModFOLDdockR | 0.464 | 0.366 | 0.191 | 0.148 |
| ModFOLDdockS | 0.394 | 0.377 | 0.177 | 0.169 |
| ModFOLDdock | 0.265 | 0.209 | 0.286 | 0.191 |
| Venclovas | 0.314 | 0.225 | 0.302 | 0.341 |

**Supplementary Table S15.**
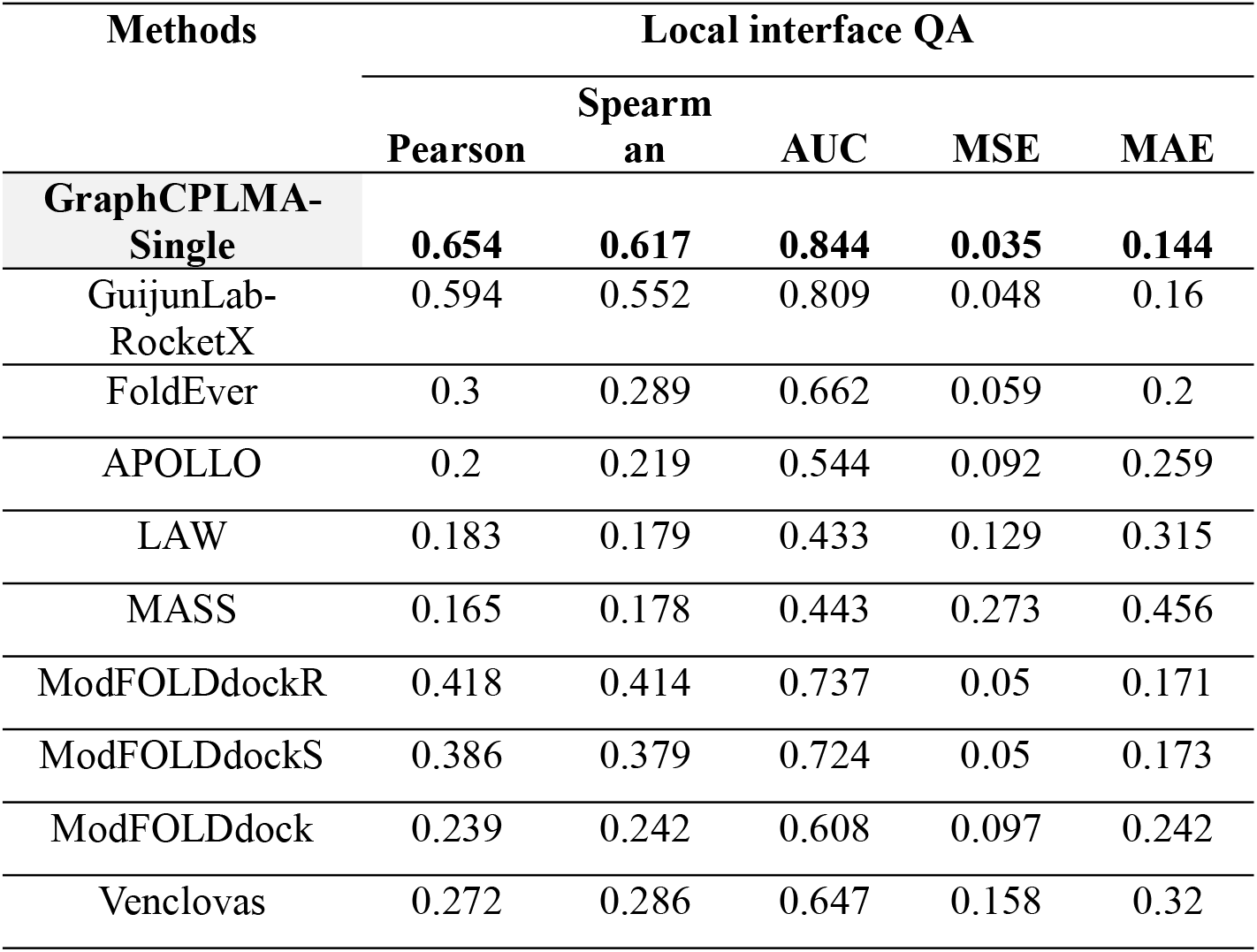
Comparison of GraphCPLMQA-Single with other methods on models of CASP15

| Methods | Local interface QA |  |  |  |  |
| --- | --- | --- | --- | --- | --- |
|  | Pearson | Spearman | AUC | MSE | MAE |
| **GraphCPLMA-Single** | **0.654** | **0.617** | **0.844** | **0.035** | **0.144** |
| GuijunLab-RocketX | 0.594 | 0.552 | 0.809 | 0.048 | 0.16 |
| FoldEver | 0.3 | 0.289 | 0.662 | 0.059 | 0.2 |
| APOLLO | 0.2 | 0.219 | 0.544 | 0.092 | 0.259 |
| LAW | 0.183 | 0.179 | 0.433 | 0.129 | 0.315 |
| MASS | 0.165 | 0.178 | 0.443 | 0.273 | 0.456 |
| ModFOLDdockR | 0.418 | 0.414 | 0.737 | 0.05 | 0.171 |
| ModFOLDdockS | 0.386 | 0.379 | 0.724 | 0.05 | 0.173 |
| ModFOLDdock | 0.239 | 0.242 | 0.608 | 0.097 | 0.242 |
| Venclovas | 0.272 | 0.286 | 0.647 | 0.158 | 0.32 |

